# Insights into regulation of C_2_ and C_4_ photosynthesis in *Amaranthaceae/Chenopodiaceae* using RNA-Seq

**DOI:** 10.1101/2021.09.14.460237

**Authors:** Christian Siadjeu, Maximilian Lauterbach, Gudrun Kadereit

## Abstract

*Amaranthaceae* (incl. *Chenopodiaceae*) show an immense diversity of C_4_ syndromes. More than 15 independent origins of C_4_ photosynthesis, partly in halophytic and/or succulent lineages, and the largest number of C_4_ species in eudicots signify the importance of this angiosperm lineage in C_4_ evolution. Here, we conduct RNA-Seq followed by comparative transcriptome analysis of three species from *Camphorosmeae* representing related clades with different photosynthetic types: *Threlkeldia diffusa* (C_3_), *Sedobassia sedoides* (C_2_), and *Bassia prostrata* (C_4_). Results show that *B. prostrata* belongs to the NADP-ME type and core genes encoding for C_4_ cycle are significantly up-regulated when compared to *Sed. sedoides* and *T. diffusa, Sedobassia sedoides* and *B. prostrata* share a number of up-regulated C_4_-related genes, however, two C_4_ transporters (DIT and TPT) are found significantly up-regulated only in *Sed. sedoides*. Combined analysis of transcription factors (TFs) of the closely related lineages (*Camphorosmeae* and *Salsoleae*) revealed that no C_3_ specific TFs is higher in C_2_ species as compared to C_4_ species, instead the C_2_ species show their own set of up-regulated TFs. Taken together, our study indicates that the hypothesis of the C_2_ photosynthesis as a proxy towards C_4_ photosynthesis is questionable in *Sed. sedoides* and more in favour of an independent evolutionary stable-state.

**Highlight:** Transcript expression profiles of C_2_ species are distinct and best explained as representing an independent evolutionary stable state.

## Introduction

C_4_ photosynthesis is a carbon-concentration mechanism, enhancing CO_2_ at the site of ribulose-1,5-bisphosphate carboxylase/oxygenase (RuBisCO). This mechanism leads to a decrease of the oxygenation reaction of RuBisCO, which in turn decreases photorespiration, because less toxic compounds resulting from the RuBisCO oxygenation reaction need to be recycled (Bauwe *et al*., 2010). C_4_ photosynthesis requires a series of biochemical, anatomical and gene regulation changes to the C_3_ photosynthesis ancestor (Gowik and Westhoff, 2011; Christin *et al*., 2013). C_4_ photosynthesis has been a major subject in life science. Since its discovery more than 50 years, its evolution is still under debate (Kadereit *et al*., 2017). The current model of C_4_ evolution relies heavily on the C_3_-C_4_ intermediate (including C_2_) photosynthetic types as evolutionary stepping stones towards C_4_ photosynthesis (Sage *et al*., 2014; Bräutigam and Gowik, 2016; Schlüter and Weber, 2020). Most C_3_-C_4_ intermediate species utilize C_2_ photosynthesis, where a photorespiratory glycine shuttle and its decarboxylation by glycine decarboxylase (GDC) concentrate CO_2_ in a bundle-sheath-like compartment (Sage *et al*., 2014). The establishment of this glycine-based CO_2_ pump and the restriction of GDC activity in the bundle sheath cells (BSCs) are considered as an important intermediate step in the evolution towards C_4_ photosynthesis (Sage *et al*., 2018). However, the absence of C_4_ relatives in lineage with C_3_-C_4_ intermediate phenotypes indicates that C_2_ photosynthesis can be an evolutionary stable stage in their own right (Monson *et al*., 1984; Edwards and Ku, 1987). On the other hand, hybrid origin of C_2_ photosynthesis was suggested because hybrids that resembled the C_3_-C_4_ intermediate using a glycine shuttle to concentrate CO_2_ anatomically was obtained by crossing *Atriplex prostrata* (C_3_) and *A. rosea* (C_4_) (Oakley *et al*., 2014).

Despite its complexity, the C_4_ pathway independently evolved in at least 61 lineages in both monocot and eudicot lineages (Sage, 2016). In eudicots, the *Amaranthaceae*/*Chenopodiaceae* alliance has the largest diversity of C_4_ syndromes with 15 independent origins of C_4_ identified, ten of which belong to *Chenopodiaceae sensu stricto* (Pyankov *et al*., 2001a; Kadereit et al., 2003; Kadereit *et al*., 2012; Kadereit *et al*., 2014; Schütze *et al*., 2003; Sage *et al*., 2007; Kadereit and Freitag, 2011; Sage, 2016). The closely related lineages *Camphorosmeae* and *Salsoleae*, belonging to the goosefoot family (*Chenopodiaceae*), are rich in C_4_ phenotypes (Kadereit and Freitag, 2011) and contain a number of C_3_-C_4_ intermediate species, including C_2_, proto-Kranz type and C_4_-like type species. Both lineages are found in steppes, semi-deserts, salt marshes and ruderal sites of Eurasia, South Africa, North America, and Australia (Akhani *et al*., 2007; Kadereit and Freitag, 2011). *Camphorosmeae* comprise subshrubs and annuals, predominantly with moderately to strongly succulent leaves with a central aqueous tissue (Kadereit *et al*., 2014). Evolutionary radiation was later in *Camphorosmeae* (early Miocene) than in *Salsoleae* (early to middle Oligocene) (Kadereit and Freitag, 2011). In *Camphorosmeae*, C_4_ photosynthesis evolved probably two times in the Miocene and different photosynthesis types are recognised on the basis of leaf anatomy with several C_4_ phenotypes (Kadereit and Freitag, 2011; Freitag and Kadereit, 2014). C_4_ photosynthesis in *Salsoleae* evolved probably multiple times and most species are C_4_ plants with terete leaves and *Salsoloid* Kranz anatomy where a continuous dual layer of chlorenchyma cells encloses the vascular and water-storage tissue (Kadereit and Freitag, 2011; Voznesenskaya *et al*., 2013; Schüssler *et al*., 2017). Therefore, these two sister groups constitute a central component allowing the investigation between and within each plant group for understanding the origin of C_2_ photosynthesis, the evolution C_4_ photosynthesis and adjustments in gene regulation leading to different photosynthesis types. Indeed, new insight into C_4_ evolution was gained from studying *Salsoleae* lineage using high-throughput sequencing methods. A transition from C_3_ cotyledons to C_4_ leaves in the *Salsoleae* lineage (*Chenopodiaceae, Salsola soda* L.) was identified (Lauterbach *et al*., 2017a). This C_3_-to-C_4_ transition is thought to be a rather exceptional phenomenon since species conducting C_4_ in all photosynthetic active tissues/organs are supposed to be the majority within this group. In addition, comparative transcriptomics revealed two proposed transporters associated with C_2_ and C_4_ photosynthesis (Lauterbach *et al*., 2017*b*). However, for *Camphorosmeae* lineage, transcriptome analysis and gene expression profile of different photosynthesis types are still lacking. Moreover, for both lineages, differential gene expression of regulatory genes (e.g. transcription factors, TFs) involved in different photosynthesis pathways remain poorly understood.

The development of complex traits is controlled by the coordination of expression of many transcription factors and signaling pathways (Monteiro and Podlaha, 2009). Thus, TFs play an important role in regulation of gene expression and are certainly responsible for the fine-tuning of the cell specific expression patterns in C_4_ photosynthesis (Aubry *et al*., 2014). The characteristic expression pattern of PHOSPHOENOLPYRUVATE CARBOXYLASE (PEPC) in C_4_ plants (i.e. high abundance in mesophyll M and low abundance in BS cells), for example, is probably established by a number of TFs from the ZINC FINGER HOMEODOMAIN (zf-HD) TF family (Windhövel *et al*., 2001). Therefore, TFs are hot candidates for a stepwise evolutionary change of complex traits such as C_4_ photosynthesis. Reviewing nine studies on potential regulators of C_4_ photosynthesis in maize, Huang and Brutnell (2016) showed that there was no TF consistently identified across these studies and suggested that consistent differential expression obtained between C_3_ and C_4_ sister lineages could be a more effective way to prioritize candidate TFs.

To fill this knowledge gap with new pieces of the puzzle, we (1) report transcriptome *de novo* assemblies and differential expression analysis between C_2_, C_3_, and C_4_ species of *Camphorosmeae* (*Amaranthaceae*) using RNA-Seq, and (2) assess transcriptional regulator elements involved in C_3_, C_2_ and C_4_ photosynthesis in the *Amaranthaceae*/*Chenopodiaceae* complex. In this latter objective, we merged transcriptome data generated in this study with the publicly available transcriptome data of C_2_, C_3_, and C_4_ species of the sister tribe *Salsoleae*.

## Materials and Methods

### Plant material

Plants of three *Camphorosmeae* species (*Bassia prostrata* (L.) Beck (C_4_), *Sedobassia sedoides* (Schrad.) Freitag & G. Kadereit (C_2_), and *Threlkeldia diffusa* R.Br. (C_3_) (Fig. 1, for voucher information see Supplementary Table S1) were grown from seeds in custom mixed potting soil in a glasshouse at Botanic Garden, Johannes Gutenberg-University Mainz, Germany with an additional light intensity of ca. 300 µmol m-2 s-1. Leaf samples of the three species were harvested between 16th April and 16th May 2014 between 10:30 am and 13:00 pm, immediately frozen in liquid nitrogen and stored at -80 °C for RNA extraction.

**Fig. 1.**
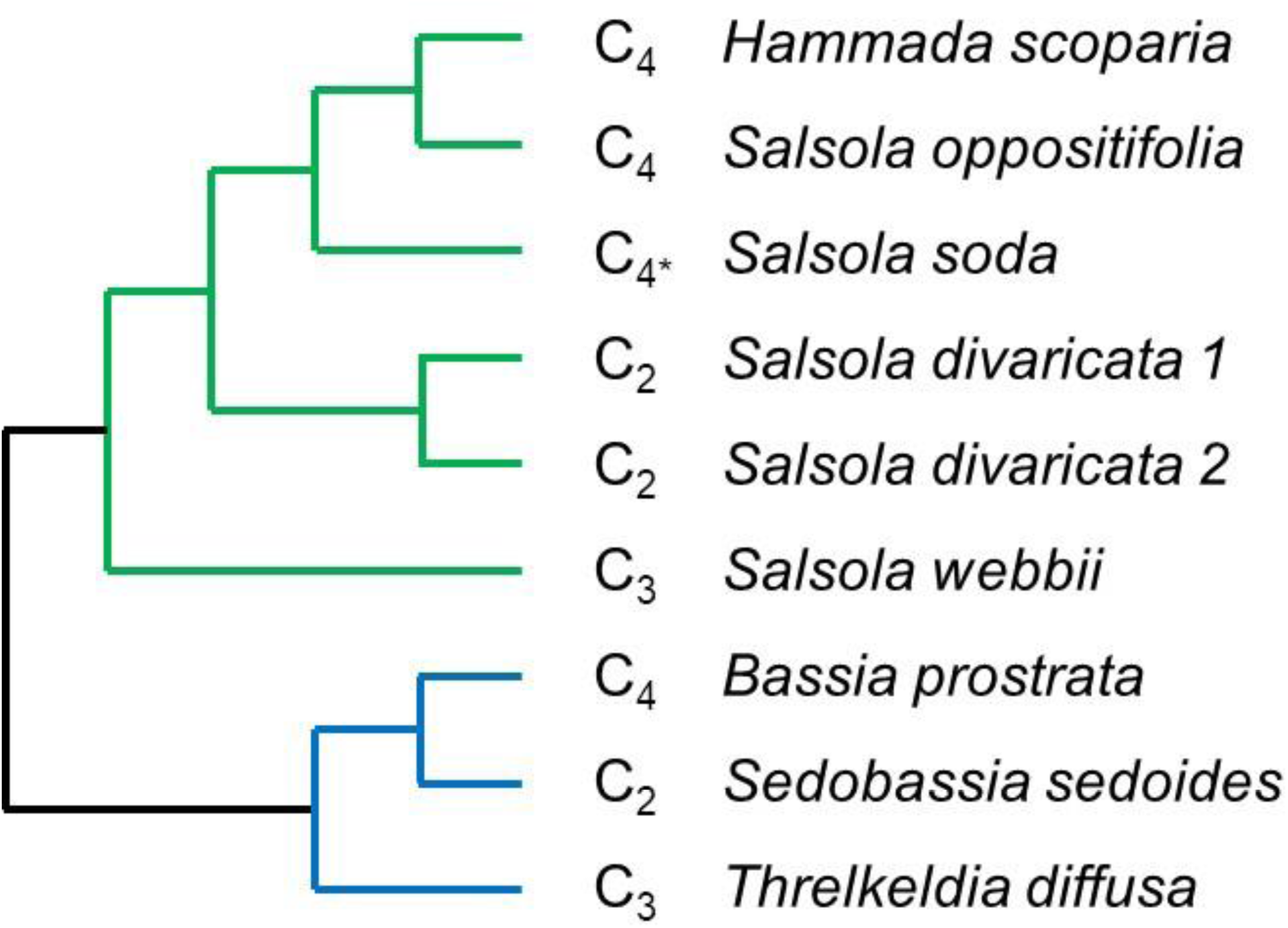
Phylogenetic relationships between species of the current study. The photosynthetic type is indicated. C_4_*, species with C_4_ photosynthesis in leaves/assimilating shoots but C_3_ in cotyledons. Green colour represents *Salsoleae*, blue *Camphorosmeae*.

### RNA isolation and sequencing

Total RNA extraction, library preparation and mRNA sequencing were performed as described in Lauterbach et al. (2017a, b). Total RNA was extracted from 16-55 mg leaf tissue of *B. prostrata, Sed. sedoides*, and *T. diffusa*. Sequencing of 101 bp single-end reads was performed on an Illumina HiSeq2000 platform. For each species, three individuals were sequenced (i.e. biological triplicates). Sequencing reads of these three species are available under study accession PRJEB36559.

### Data access

RNA-Seq data of the five *Salsoleae* species were retrieved from Lauterbach *et al*. (2017 a, b; study accession numbers PRJNA321979 and PRJEB22023) (Fig. 1, Supplementary Table S1). These data comprise: cotyledons, and first and second leaf pair of *Salsola soda* (C_3_/C_4_), cotyledons and leaves of *Salsola divaricata* population 184 (C_2_, Pop-184), *Salsola divaricata* population 198 (C_2_, Pop-198), and *Salsola oppositifolia* (C_4_); leaves of *Salsola webbii* (C_3_); and assimilating shoots of *Hammada scoparia* (C_4_). For all these samples, triplicates per species and organ were available (Lauterbach *et al*., 2017 a, b).

### RNA-Seq data processing

Single-end sequencing reads were checked for quality using FASTQC tool (www.bioinformatics.babraham.ac.uk/projects/fastqc/), and filtered and trimmed using Trimmomatic v.0.38 (Bolger *et al*., 2014). For each species, *de novo* assembly was conducted using quality-filtered reads of all replicates of leaves and, where present, cotyledons of the respective species with default parameters in Trinity v.2.1.1 (Grabherr *et al*., 2013). Quality of assemblies were assessed with BUSCO v.3.0 (Benchmarking Universal Single-Copy Orthologs, Simão *et al*. (2015)) using the ‘Eudicotyledons odb10’ dataset (Kriventseva *et al*., 2019). Number of contigs of *de novo* assemblies were reduced by clustering via CD-HIT-EST v.4.7 (Li and Godzik, 2006; Fu *et al*., 2012) and only contigs with an open reading frame were included in the downstream analysis, which was conducted with TransDecoder v.5.3.0 (github.com/TransDecoder/TransDecoder) followed by another round of CD-HIT-EST. Orthology assignment between the nine *de novo* assemblies were done by conditional reciprocal best (crb) BLAST v.0.6.9 (Aubry *et al*., 2014) run locally using protein-coding sequences of *Beta vulgaris* (version ‘BeetSet-2’, Dohm *et al*., 2014) as reference. Only contigs were included in downstream analyses that had ortholog assignments between at least six of the eight species. Besides ‘BeetSet- 2’ from *Beta vulgaris*, contigs were annotated using *Arabidopsis* (TAIR10). Reads of each of the replicates were separately mapped against these reduced data sets via bowtie2 v.2.3.4.1 (Langmead and Salzberg, 2012). Re-formatting and final extraction of read counts (excluding supplementary alignments) were done in Samtools v.1.3 (Li *et al*., 2009).

### Differential gene expression analysis

Read counts were normalised into transcripts per million (TPM) and used for differential gene expression analysis. Here, pairwise comparison between all nine species were statistically evaluated using edgeR (Robinson *et al*., 2010) in R (R Core Team, 2018). Hierarchical clustering using Pearson correlation and principal component analysis of log_2_ transformed reads counts were done with Multiexperiment Viewer (MeV) v.4.9 (http://mev.tm4.org/). Co-expressed gene clusters of (1) all expressed transcripts and (2) transcripts annotated as transcription factors were done with Clust v.1.10.7 (Abu-Jamous and Kelly, 2018). Pathways were defined in MapMan4 categories (Schwacke *et al*., 2019) with the additional category ‘C_4_’. To identify TFs putatively involved in the formation/regulation of C2 and C4 photosynthesis, the 1163 annotated TFs from *Beta vulgaris* (version ‘BeetSet-2’, Dohm *et al*., 2014) from The Plant Transcription Factor Database v.5.0 (PlantTFDB; Jin *et al*., 2014, 2017) were used. Here, two different datasets were combined (1: leaf transcriptome data of the three *Camphorosmeae* species *T. diffusa* (C3), *Sed. sedoides* (C2), and *B. prostrata* (C4); 2: leaf transcriptome data of the six Salsoleae species *Sal. webbii* (C3), *Sal. divaricata* Pop-184 (C2), *Sal. divaricata* Pop-198 (C2), *H. scoparia* (C4), *Sal. oppositifolia* (C4), and *Sal. soda* (C4); for study accession numbers see above) and transcribed TFs grouped using Clust v.1.10.7 (Abu-Jamous & Kelly, 2018) and grouping all samples based on the photosynthetic type into the three conditions C3, C2, and C4. VENNY v. 2.1 (https://bioinfogp.cnb.csic.es/tools/venny/) were deployed to find intersected TFs across all pairwise comparisons.

## Results

### Descriptive statistics of RNA data and de novo assembly

Between 27.4 and 37.9 million reads remained after quality filtering (98.68-99.06%) and were *de novo* assembled for each of the three species (Supplementary Dataset S1). Reduction of contigs by clustering resulted in 26,842 (*Sed. sedoides*, C_2_), 33,1653 (*T. diffusa*, C_3_), and 34,278 (*B. prostrata*, C_4_) contigs with an open reading frame. The number of BUSCOs genes recovered was 88.1, 88.6 and 90.9% for *T. diffusa, B. prostrata* and *Sed. sedoides*, respectively (Supplementary Table S2).

### Differential expression genes (DEGs) within Camphorosmeae

In total, 10,513 transcripts were expressed in at least six of the eight species and downstream analyses focused on this dataset (Supplementary Dataset S2). Principal component analysis of log_2_ transformed read counts (normalized to TPM) showed that replicates of each species in *Camphorosmeae* were very similar while different species were clearly distinct from each other (Fig. 2). The first principal component explained 51.47% of the total variation and *Sed. sedoides* (C_2_) was positioned somewhere in between *B. prostrata* (C_4_) and *T. diffusa* (C_3_). This result was similar to what was found in *Salsoleae* (Lauterbach *et al*., 2017b) in terms of that the C_3_-C_4_ intermediate species was positioned in between the C_3_ and the C_4_ species. The second component, explaining 42.51% of total variation, separated *Sed. sedoides* from the other two species.

**Fig. 2.**
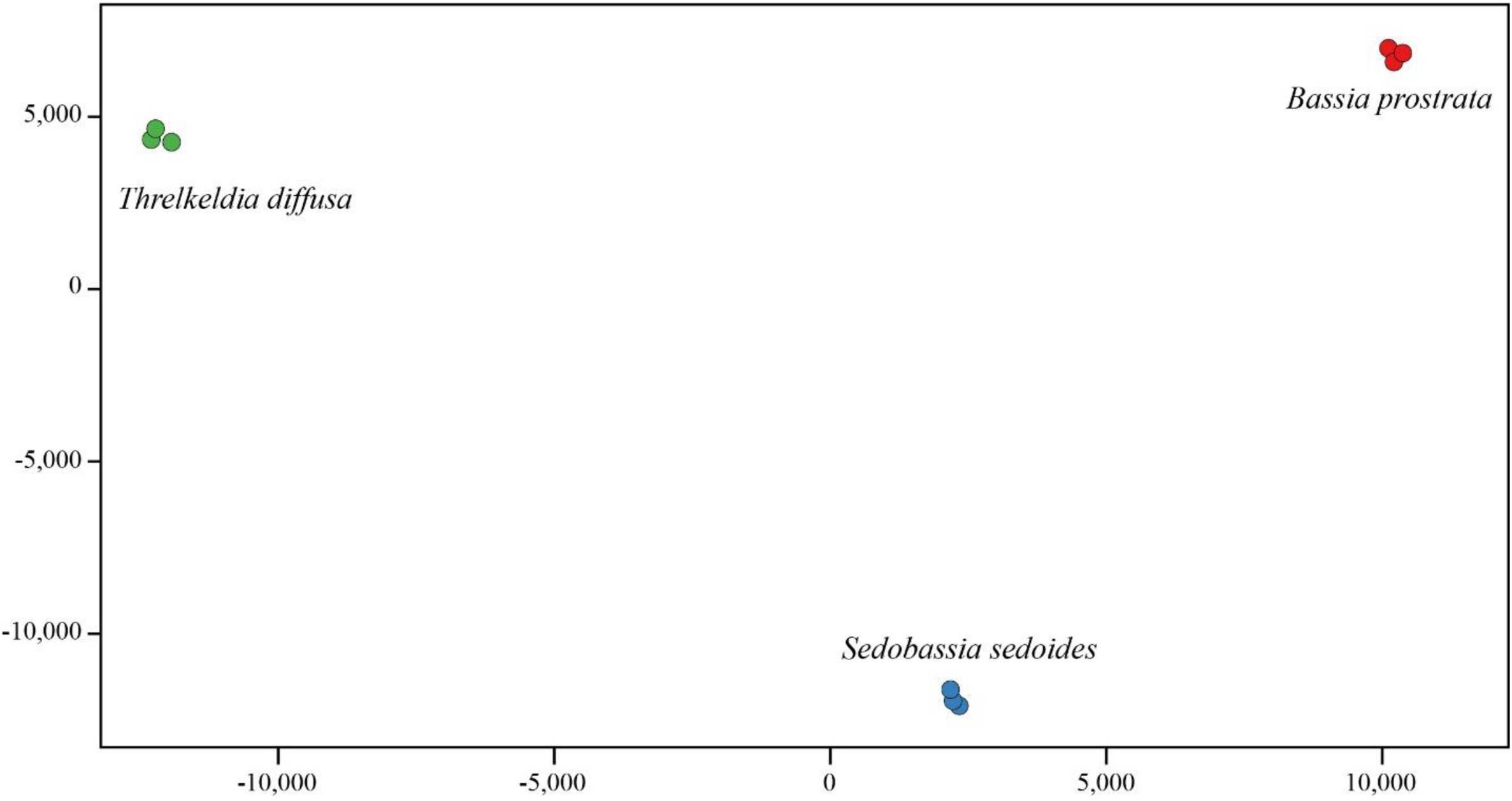
Principal component analysis of log_2_-transformed reads. The first (X-axis) and the second (Y-axis) components are shown, which explain 51.47% and 42.51% of the total variation, respectively.

### Functional classification and enrichment of DEGs within the Camphorosmeae

In all three species, transcriptional investment, defined as percentage of all read counts of transcripts (normalized to TPM) belonging to a particular MapMan category, was highest in the MapMan category ‘Not assigned’ (23.10-27.54%) (Fig. 3, Supplementary Dataset S3). MapMan category ‘Photosynthesis’ was second highest in all three species, however, the amount differed between species from 11.24% in *B. prostrata* (C_4_), 15.03% in *T. diffusa* (C_3_), up to 21.58% in *Sed. sedoides* (C_2_) (Fig. 3, Supplementary Dataset S3). In general, high transcriptional investment in the category ‘Photosynthesis’ in *Sed. sedoides* (C_2_) was caused by higher transcription of many genes of the sub-category ‘Calvin cycle’. However, transcription of a gene encoding RUBISCO ACTIVASE in *Arabidopsis* (AT2G39730) was the main driver of the difference among species (*Sed. sedoides* compared to *T. diffusa*: log2 fold change of 2.09; *Sed. sedoides* compared to *B. prostrata*: log2 fold change of 3.72; Supplementary Dataset S2). Transcripts belonging to the categories ‘Protein biosynthesis’ (5.93-8.27%), ‘Protein degradation’ (4.45-5.98%), and ‘Protein modification’ (3.03-4.09%) were also highly abundant in all three species. In *B. prostrata*, the category ‘C_4_’ was higher (transcriptional investment of 4.54%) compared to in *Sed. sedoides* (1.09%) and *T. diffusa* (0.58%; Fig. 3, Supplementary Dataset S3). ‘Photorespiration’ was about twice as high in *Sed. sedoides* and *T. diffusa* (2.59% and 2.21%, respectively) compared to *B. prostrata* (1.14%). This difference was caused by higher transcription of most genes of the category ‘photorespiration’ rather than few ones (Supplementary Dataset S3). The categories with the lowest transcriptional investment in all three species were ‘DNA damage response’ (0.10- 0.14%), ‘Polyamine metabolism’ (0.16-0.29%), and ‘Multi-process regulation’ (0.18-0.34%).

**Fig. 3.**
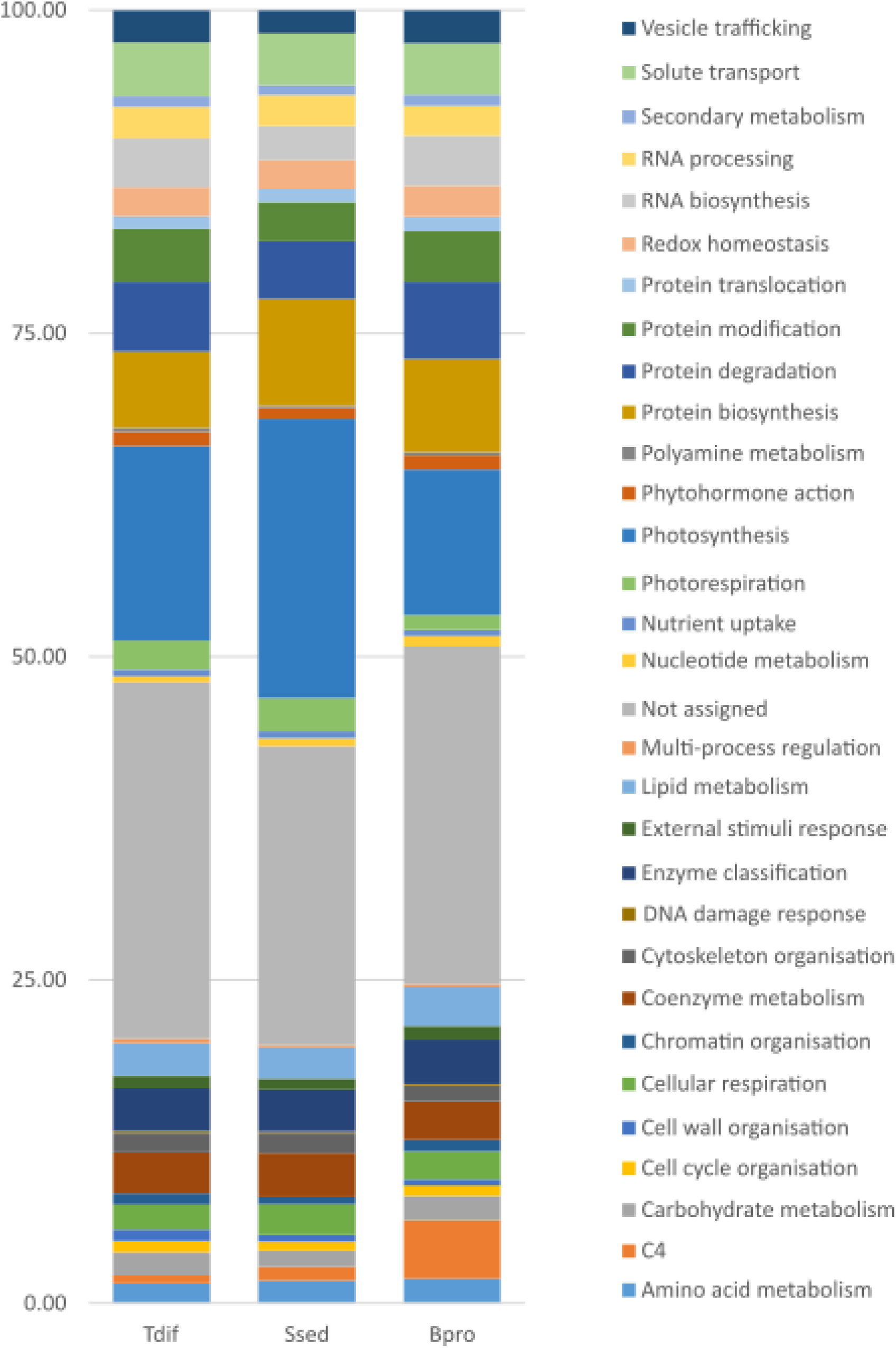
Distribution of transcriptional investment defined as the percentage of all transcripts belonging to a particular MapMan4 category (Schwacke *et al*., 2019) and the additional categories ‘C_4_’.

### Differential expression of C_4_-related genes in C_3_, C_2_ and C_4_ Camphorosmeae species

Like observed in other C4 species, most genes encoding for proteins involved in C4 photosynthesis were significantly up-regulated in the C4 species *B. prostrata* (C_4_) when compared to *Sed. sedoides* (C2) and *T. diffusa* (C3) (Table 1, Supplementary Dataset S2-S4). Out of 18 C_4_- related transcripts, 16 and 13 transcripts encoding known C_4_ cycle proteins were significantly up- regulated (P < 0.001) in *B. prostrata* (C_4_) compared to *T. diffusa* (C_3_) and *Sed. sedoides* (C2) respectively. ALANINE AMINOTRANSFERASE (AlaAT, Log_2_FC = 5.17) was the most abundant followed by PYRUVATE ORTHOPHOSPHATE DIKINASE (PPdK, Log_2_FC=4.63), PHOSPHOENOLPYRUVATE CARBOXYLASE (PEPC, Log_2_FC =4.12), BILE ACID:SODIUM SYMPORTER FAMILY PROTEIN 2 (BASS2, Log_2_FC = 3.97), NADP-malic enzyme (NADP-ME Log_2_FC = 3.86), PEP/phosphate translocator (PPT, Log_2_FC = 3.61) in *B. prostrata* compared to *T. diffusa*. Conversely, BASS2 Log_2_FC = 3.51, PPdK Log_2_FC = 3.37, PHOSPHATE TRANSPORTER 1 (PHT1, Log_2_FC = 3.16), PEPC (Log_2_FC = 2.95), AlaAT (Log_2_FC = 2.61) were the top five of highly up-regulated genes in *B. prostrata* (C_4_) when compared to *Sed. sedoides* (C_2_). However, all C_4_-related genes except TRANSPLANTA (TPT), a Carbonic anhydrase (CA) isoform (Bv8_194450_rkme.t1) and a chloroplast dicarboxylate transporter isoform DiT (Bv4_072630_xjai.t1) significantly up-regulated in *B. prostrata* (C_4_) when compared to *T. diffusa* (C_3_) were significantly up-regulated in *B. prostrata* (C_4_) as compared to *Sed. sedoides* (C_2_). On the other hand, transporters [TPT (Bv8_194450_rkme.t1), DiT (Bv4_072630_xjai.t1)] and a CA isoform (Bv8_194450_rkme.t1) were significantly up-regulated in *Sed. sedoides* (C_2_) compared to *B. prostrata* (C_4_) (Supplementary Dataset S2). Interesting, PHT1 (Bv3_049110_qgnh.t1) significantly up-regulated in *B. prostrata* (C_4_) when compared to *Sed. sedoides* (C_2_) (Table 1) was significantly up-regulated in *T. diffusa* (C_3_) as compared to *Sed. sedoides* (C_2_) (Supplementary Dataset S2).

**Table 1.** Differential expression of C_4_-related enzymes in leaves of *Camphorosmeae* species. *T. diffusa* (C_3_), *Sed. sedoides* (C_2_), *B. prostrata* (C_4_)

| Transcripts | genes | C<sub>2</sub> vs C<sub>4</sub>* |  | C<sub>3</sub> vs C<sub>4</sub>* |  | C<sub>3</sub> vs C<sub>2</sub>* |  |
| --- | --- | --- | --- | --- | --- | --- | --- |
|  |  | Log2FC | Padj | Log2FC | P | Log2FC | Padj |
| Bv_006710_gkqg.t1 | MDH | 1.29 | 1.01E-10 | 2.84 | 9.92E-42 | 1.55 | 1.74E-14 |
| Bv1_004490_tyfq.t1 | PHT4 | 1.57 | 1.56E-11 | 2.89 | 1.72E-31 | 1.32 | 7.56E-08 |
| Bv1_013550_fjqs.t1 | PPdK | 3.37 | 2.32E-54 | 4.63 | 2.78E-90 | 1.26 | 8.92E-10 |
| Bv2_031080_twkf.t1 | AspAt | 1.32 | 2.84E-06 | 2.15 | 1.44E-13 | 0.82 | 0.003667 |
| Bv3_049110_qgnh.t1 | PHT1 | 3.16 | 8.05E-08 | - | - | - | - |
| Bv4_072630_xjai.t1 | DiT | - | - | 1.30 | 4.88E-06 | 2.61 | 3.46E-19 |
| Bv5_117240_yhsk.t1 | PPT | 1.94 | 7.13E-13 | 3.61 | 5.61E-35 | 1.67 | 2.18E-09 |
| Bv6_135140_uyxu.t1 | Asn Synthetase | 2.56 | 5.60E-07 | 2.42 | 1.93E-06 | - | - |
| Bv6_148840_uffy.t1 | CA | 3.80 | 6.66E-38 | 3.81 | 4.08E-38 | - | - |
| Bv7_169130_kwer.t1 | AlaAT | 2.61 | 4.89E-17 | 5.17 | 4.74E-48 | 2.56 | 2.34E-15 |
| Bv8_182550_kstq.t1 | TPT | - | - | 1.48 | 1.20E-10 | 2.29 | 2.71E-22 |
| Bv8_194450_rkme.t1 | CA | - | - | - | - | 1.48 | 0.000263 |
| Bv8_195530_sxjq.t1 | BASS2 | 3.51 | 5.00E-68 | 3.97 | 6.26E-83 | 0.46 | 0.018625 |
| Bv8_200290_ujgk.t1 | TPT |  |  | 3.25 | 1.25E-05 | 1.97 | 0.005513 |
| Bv9_209750_xeaz.t1 | PEPC | 2.95 | 3.29E-17 | 4.12 | 6.98E-29 | 1.18 | 0.000429 |
| Bv9_215520_prze.t1 | DiT | 1.48 | 1.23E-08 | 2.63 | 5.15E-22 | 1.16 | 2.27E-05 |
| Bv9_224000_xpgi.t1 | NHD | 2.48 | 1.17E-21 | 2.68 | 7.04E-25 | - | - |
| Bv9_224840_zmjw.t1 | NADP-ME | 1.29 | 8.44E-08 | 3.86 | 9.54E-47 | 2.56 | 1.34E-23 |

- not significantly expressed Pvalue > 0.05

Twelve of C_4_-related enzymes except ASPARAGINE SYNTHETASE (Asn Synthetase), SODIUM:HYDROGEN ANTIPORT (NHD) and a CARBONIC ANHYDRASE (CA) isoform (Bv6_148840_uffy.t1) were significantly up-regulated in *Sed. sedoides* (C_2_) as compared to *T. diffusa* (C_3_) including typical C_4_ enzymes like PEPC, NADP-ME, PPdK, PHT4, Ala-AT, ASPARTATE AMINOTRANSFERASE (Asp-AT), as well as C_4_ associated transporters such as BASS2 and DiT.

### Differential expression of key photorespiration genes in C_3_, C_2_ and C_4_ Camphorosmeae species

Out of 14 transcripts associated with photorespiratory enzymes, 12 were annotated and assigned (Table 2). All these 12 photorespiratory transcripts were significantly up-regulated in *Sed. sedoides* (C_2_) as compared to *B. prostrata* (C_4_) including the core photorespiratory enzymes GLYCINE DECARBOXYLASE (GDC, T-, H-, P-, L-), GLUTAMATE:GLYOXYLATE AMINOTRANSFERASE (GGT) and SERINE HYDROXYMETHYLTRANSFERASE (SHMT) among the top ten. In *Sed. sedoides* (C_2_), GGT, two SHMTs, GDC-T, GLYCOLATE OXIDASE (GOX), GDC-H, GLYCERATE 3-KINASE (GLYK), PHOSPHOGLYCOLATE PHOSPHATASE (PGP) were significantly up-regulated when compared to *T. diffusa* (C_3_). Only one SHMT isoform (Bv6_143730_mggd.t1) was significantly upregulated in *B. prostrata* (C_4_) when compared to *Sed. sedoides* (C_2_) (Supplementary Dataset S2). In *T. diffusa* (C_3_), only one gene GDC-P was significantly up-regulated as compared to *Sed. sedoides* (C_2_) (Supplementary Dataset S2). All photorespiratory genes except GLYK significantly expressed in *Sed. sedoides* (C_2_) as compared to *B. prostrata* (C_4_) were significantly up-regulated in *T. diffusa* (C_3_) when compared to *B. prostrata* (C_4_).

**Table 2.**
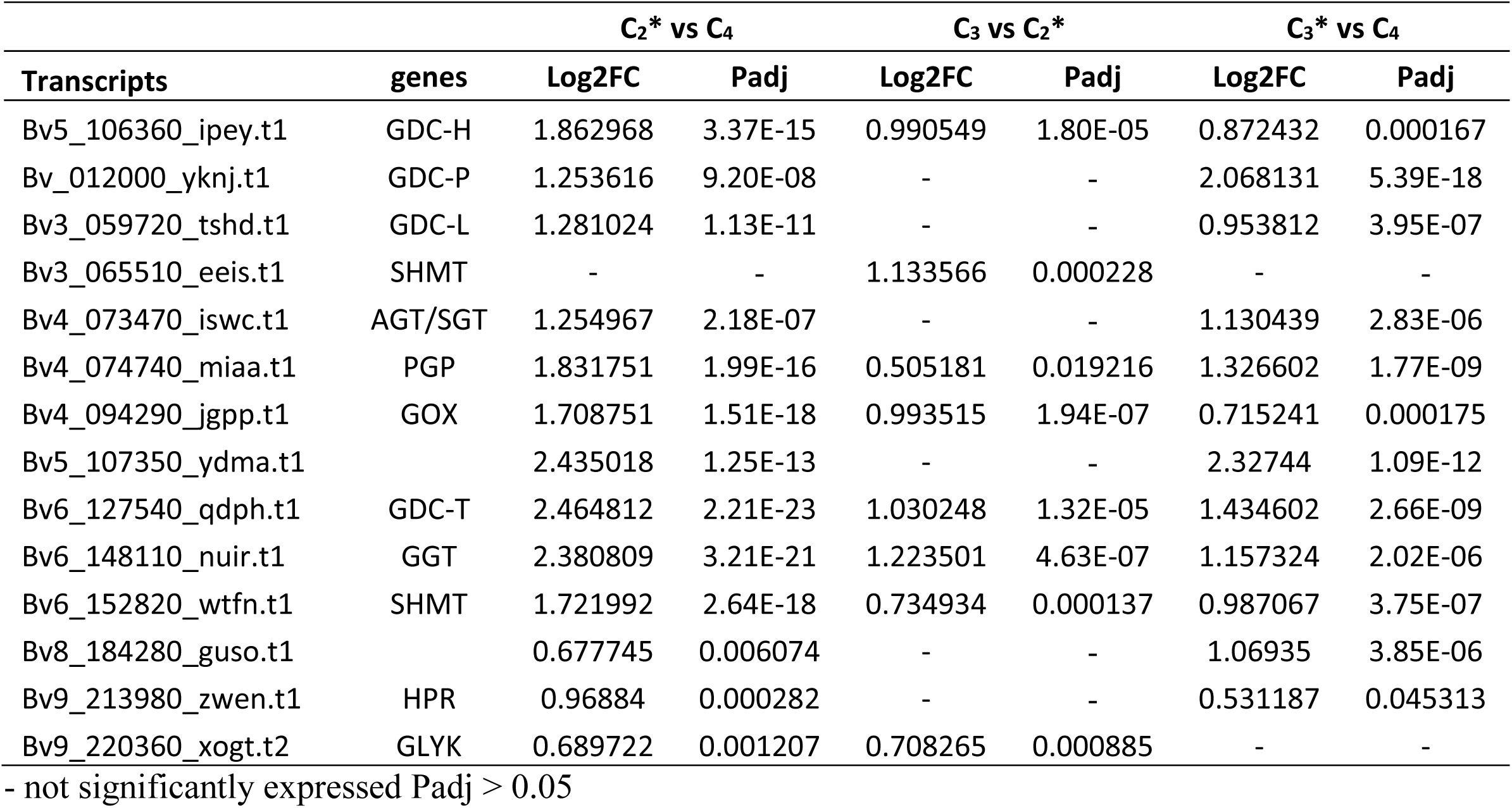
Differential expression of photorespiratory transcripts in leaves between *Camphorosmeae* species. *T. diffusa* (C_3_), *Sed. sedoides* (C_2_), *B. prostrata* (C_4_)

### Regulatory elements in C_3_, C_2_, and C4 species of the Amaranthaceae/Chenopodiaceae alliance

To identify regulatory genes putatively involved in the formation/regulation of C2 and C4 photosynthesis, expression patterns of TFs from The Plant Transcription Factor Database v.5.0 (PlantTFDB; Jin *et al*., 2014, 2017) using 1163 annotated TFs from *Beta vulgaris* (version ‘BeetSet-2’, Dohm *et al*., 2014) were investigated. From the whole set of TFs included in PlantTFDB, 824 orthologous TFs were found, of which 494 TFs had orthologs in all eight species [three species from *Camphorosmeae*: *B. prostrata* (C_4_), *Sed. sedoides* (C_2_), *T. diffusa* (C_3_); five species from *Salsoleae*: *H. soparia* (C_4_), *Sal. divaricata* Pop-184 (C_2_), *Sal. divaricata* Pop-198 (C_2_), *Sal. oppositifolia* (C_4_), *Sal. soda* (C_3_/C_4_), *Sal. webbii* (C_3_)] and were further analysed using Clust. Based on the expression pattern, from the initial 494 TFs, 431 TFs were grouped into 11 clusters with between 22 and 71 genes per cluster (Cluster-C0 to Cluster-10; Fig. 4, Supplementary Fig. S6). Five of the 11 clusters were of particular interest, because these clusters included TFs that were higher abundant either in C4 species when compared to C3 and C_2_ species (Cluster-C4, Cluster-C5, and Cluster-C6) or in C3 species as compared to C4 and C_2_ species (Cluster-C0, Cluster-C9, and Cluster-C10) or in C_2_ species when compared to C_3_ and C_4_ species (Cluster-C1, Cluster-C2, Cluster-C3) (Fig. 4).

**Fig. 4.**
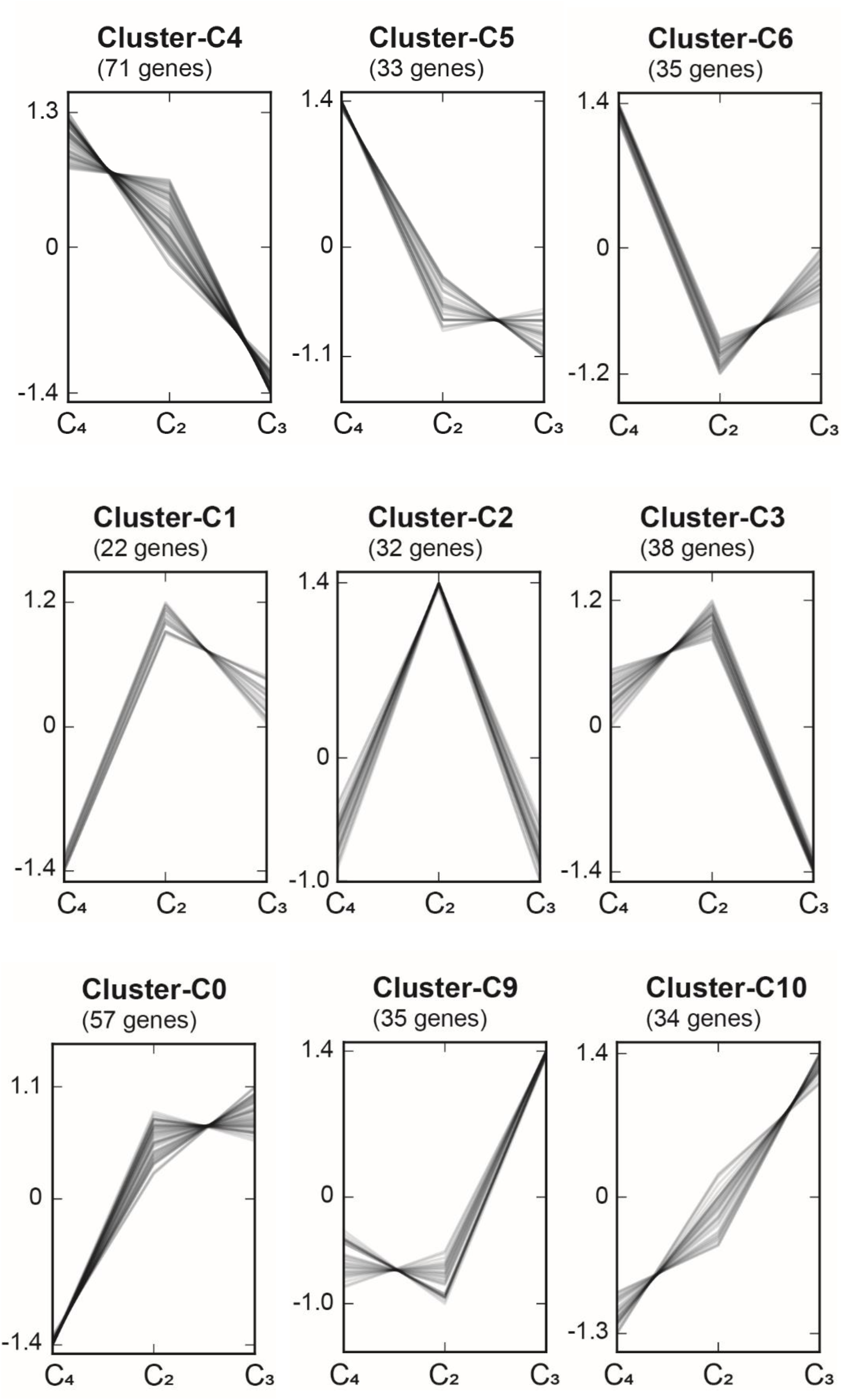
Co-expressed gene clusters were generated using Clust v.1.10.7 (Abu-Jamous & Kelly, 2018). The nine species were grouped into three different conditions ‘C_4_ species’, ‘C_2_ species’, and ‘C_3_ species’. The 11 clusters contained between 22 and 71 genes.

Cluster-C4, Cluster-C5 and Cluster-C6 consisted of 71, 33 and 35 TFs, respectively, from 41 different TF families (Fig. 4, Supplementary Dataset S5). Four TFs, all part of Cluster-C4, were present in all nine species, transcripts significantly (adjusted *P*-value ≤0.01) higher abundant in all C4 species compared to the two C3 species (Table 3). These TFs comprised BBX15 (CO-like family, AT1G25440.1), SHR (TF family GRAS), SCZ (TF family HSF), and LBD41 (TF family LBD). Cluster-C0, Cluster-C9, and Cluster-C10 respectively comprised 57, 35, and 34 TFs (Fig. 4, Supplementary Dataset S5). Here, two TFs of Cluster-10 (HSF, and NAC) and one of Cluster- 9 (HD-ZIP) were significantly abundant in the studied C3 species when compared to C4 species (Table 4).

**Table 3.**
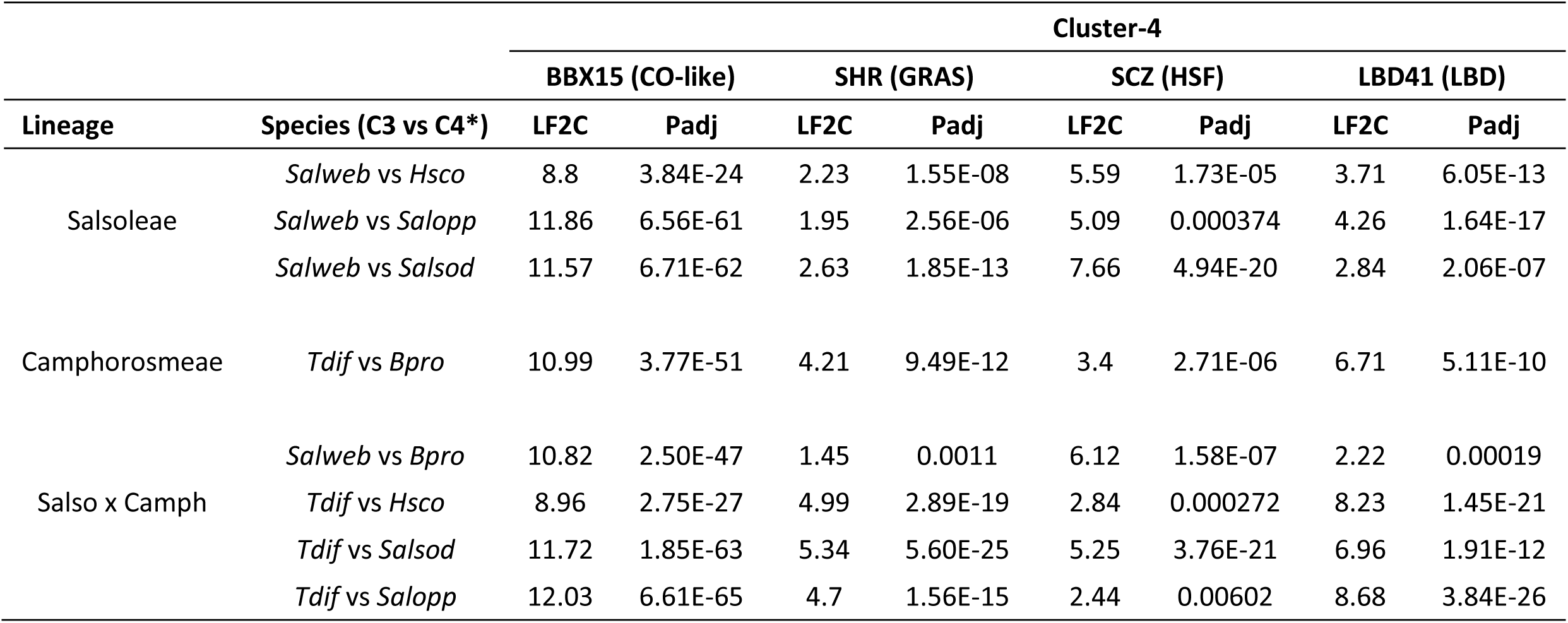
Differentially expressed C_4_-related TFs in *Amaranthaceae*/*Chenopodiaceae*

| Lineage | Species (C3 vs C4*) | Cluster-4 |  |  |  |  |  |  |  |
| --- | --- | --- | --- | --- | --- | --- | --- | --- | --- |
|  |  | BBX15 (CO-like) |  | SHR (GRAS) |  | SCZ (HSF) |  | LBD41 (LBD) |  |
|  |  | LF2C | Padj | LF2C | Padj | LF2C | Padj | LF2C | Padj |
| Salsoleae | *Salweb* vs *Hsco* | 8.8 | 3.84E-24 | 2.23 | 1.55E-08 | 5.59 | 1.73E-05 | 3.71 | 6.05E-13 |
|  | *Salweb* vs *Salopp* | 11.86 | 6.56E-61 | 1.95 | 2.56E-06 | 5.09 | 0.000374 | 4.26 | 1.64E-17 |
|  | *Salweb* vs *Salsod* | 11.57 | 6.71E-62 | 2.63 | 1.85E-13 | 7.66 | 4.94E-20 | 2.84 | 2.06E-07 |
| Camphorosmeae | *Tdif* vs *Bpro* | 10.99 | 3.77E-51 | 4.21 | 9.49E-12 | 3.4 | 2.71E-06 | 6.71 | 5.11E-10 |
| Salso x Camph | *Salweb* vs *Bpro* | 10.82 | 2.50E-47 | 1.45 | 0.0011 | 6.12 | 1.58E-07 | 2.22 | 0.00019 |
|  | *Tdif* vs *Hsco* | 8.96 | 2.75E-27 | 4.99 | 2.89E-19 | 2.84 | 0.000272 | 8.23 | 1.45E-21 |
|  | *Tdif* vs *Salsod* | 11.72 | 1.85E-63 | 5.34 | 5.60E-25 | 5.25 | 3.76E-21 | 6.96 | 1.91E-12 |
|  | *Tdif* vs *Salopp* | 12.03 | 6.61E-65 | 4.7 | 1.56E-15 | 2.44 | 0.00602 | 8.68 | 3.84E-26 |

**Table 4.** Differentially expressed C_3_-related TFs in *Amaranthaceae*/*Chenopodiaceae*.

| Lineage | Species (C4 vs C3*) | Cluster-9 |  | Cluster-10 |  |  |  |
| --- | --- | --- | --- | --- | --- | --- | --- |
|  |  | ATHB13 (HD-ZIP) |  | HSFA6B (HSF) |  | NAC083 (NAC) |  |
|  |  | LF2C | Padj | LF2C | Padj | LF2C | Padj |
| Salsoleae | *Hsco* vs *Salweb* | 9.02 | 5.45E-24 | 1.21 | 0.00017 | 0.73 | 0.00398 |
|  | *Salopp* vs *Salweb* | 1.71 | 2.77E-05 | 1.54 | 1.32E-05 | 1.42 | 4.44E-08 |
|  | *Salsod* vs *Salweb* | 1.14 | 0.00077 | 1.43 | 1.65E-06 | 0.76 | 0.0004 |
| Camphorosmeae | *Bpro* vs *Tdif* | 8.89 | 2.80E-23 | 1.7 | 1.08E-06 | 11.74 | 3.28E-82 |
| Salso. x Camph. | *Bpro* vs *Salweb* | 8.99 | 8.45E-24 | 1.91 | 5.76E-08 | 11.28 | 1.31E-72 |
|  | *Hsco* vs *Tdif* | 8.92 | 1.78E-23 | 1 | 0.00169 | 1.01 | 5.11E-05 |
|  | *Salsod* vs *Tdif* | 0.87 | 0.00956 | 1.23 | 4.09E-05 | 1.04 | 6.71E-07 |
|  | *Salopp* vs *Tdif* | 1.45 | 0.00038 | 1.34 | 0.00011 | 1.71 | 4.00E-11 |

Clustering of transcribed genes not only resulted in clusters putatively interesting for investigating C4 photosynthesis (see above), but was also able to find three clusters that included TFs higher abundant in C2 species compared to C3 and C_4_ species (Cluster-1, Cluster-C2 and Cluster-C3) (Fig. 4). Cluster-C1, Cluster-C2 and Cluster-C3 included 22, 32 and 38 TFs, respectively, of which one TF of Cluster-2 bHLH 106 (TF family bHLH) was significantly higher in the C_2_ species when compared to the C3 species and C_4_ species (Table 5, Supplementary Dataset S2-S5). To assess the integration of C_3_ and C_4_ pathways into the intermediate C_2_ pathway at the regulation level and vice versa, specific TFs of C_4_ (Cluster-C4, Cluster-C5 and Cluster-C6) and C_3_ (Cluster-C0, Cluster-C9, and Cluster-C10) pathways were assessed in the following comparisons: C_2_ species vs C_3_ species and C_2_ species vs C_4_ species. Then, specific TFs of C_2_ pathway (Cluster- C1, Cluster-C2 and Cluster-C3) specific were estimated in the pairwise comparison of C_3_ species vs. C_4_ species. Among the four TFs common to all C_4_ species, only one TF (BBX15, TF family CO-like) was significantly up-regulated in C_2_ when compared to C_3_ species (Supplementary Dataset S2-S5). Conversely, no TF of C_3_ species was highly expressed in C_2_ species when compared to C_4_ species.

**Table 5.**
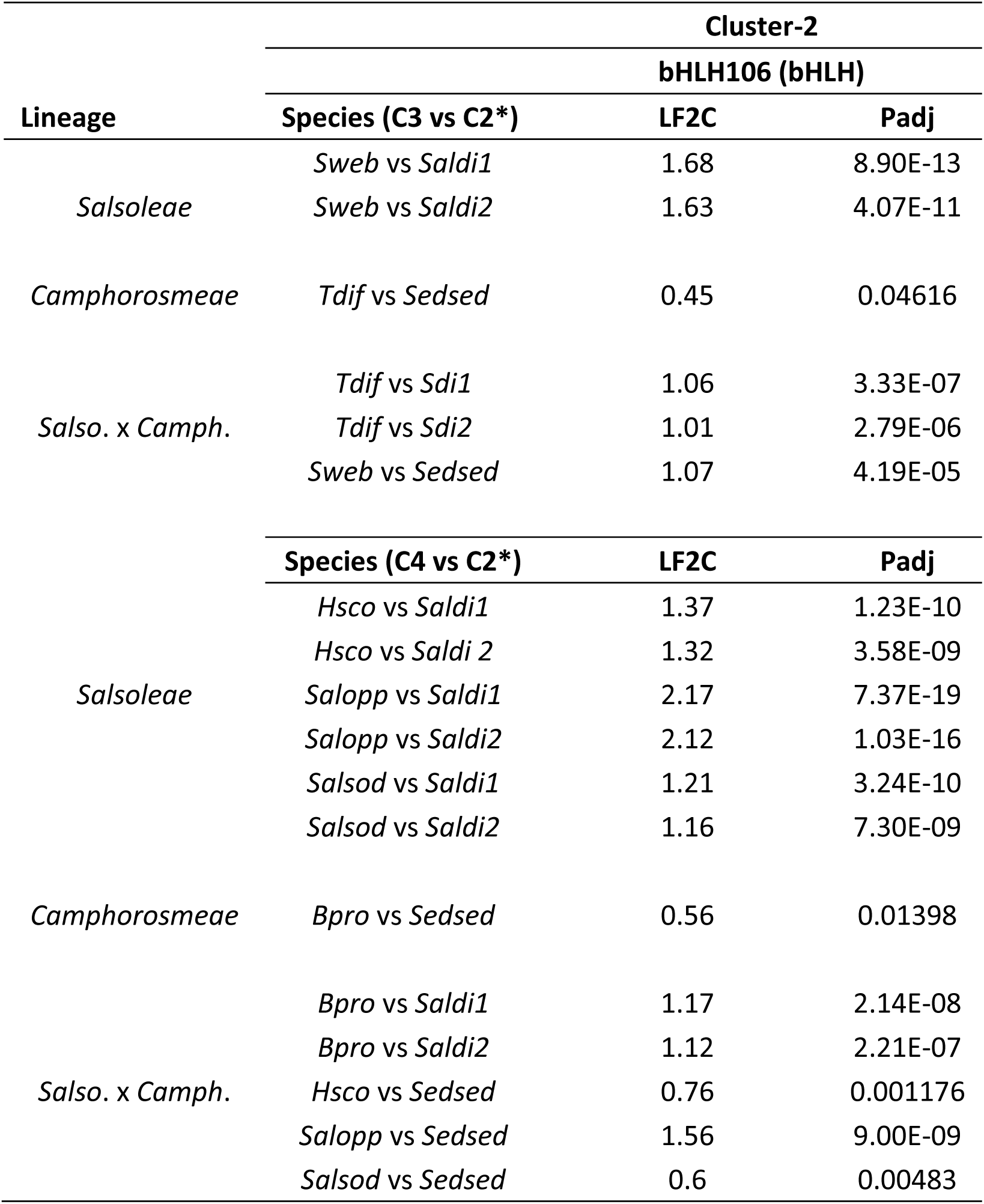
Differentially expressed C_2_-related TF in *Amaranthaceae*/*Chenopodiaceae*

## Discussion

### Transcriptome analysis in Camphorosmeae

Gene expression analysis paved the way to understand the difference between derived photosynthesis types (C_2_, C_4_) and the ancestral C_3_ photosynthesis (Gowik et al., 2011; Mallmann et al., 2014; Lauterbach et al., 2017). Much progress in understanding C_4_ and C_2_ photosynthesis was achieved by comparing differentially expressed genes of closely related species in the genus *Flaveria* (Gowik *et al*., 2011; Schulze *et al*., 2013; Mallmann *et al*., 2014; Bräutigam and Gowik, 2016; Schlüter and Weber, 2020; Sage, 2021). The goosefoot family (*Chenopodiaceae*) with a large number of C_2_ and C_4_ species that differ anatomically and ecologically from the genus *Flaveria* (*Asteraceae*) represents a good supplementary alternative to decipher the convergent evolution of C_4_ and C_2_ photosynthesis. With PCA based on gene expression, it was possible to clearly distinguish between *T. diffusa* (C_3_), representing the ancestral condition, and *Sed. sedoides* (C_2_) and *B. prostrata* (C_4_), representing derived conditions. The physiologically C_3_-C_4_ intermediate *Sed. sedoides* (C_2_) was positioned in a triangle with *T. diffusa* (C_3_) and *B. prostrata* (C_4_) in terms of transcript variation. This result was similar to what was found in *Salsoleae* (Lauterbach *et al*., 2017b). The first three components explained about 76% of the total variation, which was slightly higher than the 73% reported in *Salsoleae* (Lauterbach et al., 2017b). Similar to *Salsoleae*, in *Camphorosmeae*, the three different photosynthesis types predominantly structure the gene expression pattern in assimilating tissue. Indeed, *T. diffusa* (C_3_), *Sed. sedoides* (C_2_), and *B. prostrata* (C_4_) differ in leaf anatomy with respect to the type of photosynthesis: *Neokochia* type (C_3_ with an undifferentiated chlorenchyma of several layers), *Sedobassia* type (C_2_ with kranz-like cells near peripheral vascular bundles), *Bassia prostrata* type (C_4_ with the chlorenchyma differentiated in an outer mesophyll and inner kranz-layer) (Freitag and Kadereit, 2014). In contrast, the first three PCA components in a comparable study of *Flaveria* explained only 27% (Mallmann et al., 2014). This difference could be due to the younger evolutionary age or other confounding factors affecting gene expression in the *Flaveria* study as suggested by Lauterbach et al. (2017 b).

### C_4_ key enzymes in C_4_ and C_2_ Camphorosmeae species

Analyses of differential gene expression between C_3_ and C_4_ species of *Camphorosmeae* showed that core C_4_ cycle proteins were highly abundant in *B. prostrata* (C_4_). Similar results were found in *Cleome* (Bräutigam *et al*., 2011), *Flaveria* (Gowik *et al*., 2011; Mallmann et al., 2014) and *Salsoleae* (Lauterbach et al., 2017 b). Traditionally, three biochemical subtypes of C_4_ photosynthesis are classified according to the predominant type of decarboxylation releasing CO_2_ around RUBisCo in the BSCs: NAD-ME, NADP-ME, and PEP-CK. However, PEPCK should be considered as a supplementary subtype to either NAD-ME or NADP-ME (Wang *et al*., 2014). Significant expression of NADP-ME indicates that *B. prostrata* (C_4_) uses a NADP-ME type C_4_ cycle. Asparagine synthetase (ASN) and NHD were found significantly expressed and up- regulated only in *B. prostrata* (C_4_) as compared to *T. diffusa* (C_3_) and *Sed. sedoides* (C_2_). ASN was reported up-regulated in C_4_ species *Gynandropsis gynandra* when compared to closely related C_3_ species *Tarenaya hassleriana* (*Cleomaceae*), as well as in C_4_ leaves of *Sal. soda* when compared to its C_3_ cotyledones (Lauterbach et al., 2017 a). On the other hand, NHD was found up-regulated in C_4_ species compared to C_3_ and C_2_ species of *Flaveria* (Mallmann et al., 2014). Moreover, the top three of highly expressed C_4_ enzymes in *B. prostrata* (C_4_) as compared to *T. diffusa* (C_3_) were Ala-AT, PPDK, and BASS2. ASN is involved in ammonium metabolism and asparagine in nitrogen transport (Lauterbach et al., 2017 a). Achievement of the C_4_ cycle requires the transport of pyruvate to the mesophyll cell (MC) for regeneration of PEP. While Ala-AT plays an important role in pyruvate generation, PPDK intervenes in the regeneration of PEP. Pyruvate transport is mediated by the BASS2/NHD transport system (Furumoto *et al*., 2011). Taken all together, this indicates not only a possible functional connection between nitrogen metabolism and the switch from C_3_ to C_4_ pathway as suggested by Lauterbach *et al*. (2017b) and Mallmann et al. (2014), but also the capacity to shuttle pyruvate from the BS plastid. In this regard, to ensure the regeneration of CO_2_ acceptor (PEP), and therefore to maintain the C_4_ pathway.

Eleven C_4_-related genes were found significantly up-regulated in *Sed. sedoides* (C_2_) as compared to *T. diffusa* (C_3_) including for example PEPC, NADP-ME, PPdK, PHT4. Up-regulation of C_4_ typical enzymes such as PEPC, NADP-ME, PPdK, PPT was also reported in the C_2_ species when compared to the C_3_ species in studies of *Flaveria* and *Salsoleae* (Mallmann et al., 2014, Lauterbach et al., 2017b). This result suggests that genes associated with the C_4_ cycle are present in *Sed. sedoides* (C_2_) and play an important role in the C_2_ metabolism. A DiT isoform (Bv4_072630_xjai.t1), NADP-ME and Ala-AT were the three most up-regulated C_4_ enzymes in *Sed. sedoides* (C_2_) as compared to *T. diffusa* (C_3_). Moreover, we found two transporters (TPT and DIT) up-regulated in *Sed. sedoides* (C_2_) when compared to *B. prostrata* (C_4_). These transporters were found in four experiments inconsistently high expressed in C_2_ species when compared to C_4_ species in *Flaveria* (Mallman et al., 2014). DIT is a putative plastidial dicarboxylate transporter and the chloroplast envelope triose-phosphate/phosphate translocator (TPT) is responsible for carbohydrate export during photosynthesis (Walters *et al*., 2004). Based on simulated data, it was shown that a high TPT capacity is required to obtain high assimilation rates and to decrease the CO_2_ leakage from BSCs to MCs (Wang et al., 2014). The most likely reason for up-regulation of these genes is their involvement in decreasing the CO_2_ leakage from the Kranz-like cells back to the MCs due to the presence of RuBisCo, which is not the case for C_4_ plants. This explains the low CO_2_ compensation observed in C_2_ species (Sage *et al*., 2013). Thus, C_2_ plants up-regulate a distinct set of C_4_ enzymes to handle constraints related to the C_2_ pathway and not an entirely congruent set. This does not support their interpretation as an intermediate state towards C_4_ photosynthesis but is more in line with their interpretation as an independent evolutionary stable state (Lundgren, 2020 and refs. therein).

### Photorespiratory enzymes in C_4_ and C_2_ Camphorosmeae species

Transcripts associated with photorespiration were about twice as high in C_3_-C_4_ intermediate (*Sed. sedoides*) and *T. diffusa* (C_3_) compared to *B. prostrata* (C_4_). Likewise, we found key photorespiration enzymes were differentially expressed and up-regulated in the C_2_ species (*Sed. sedoides*) when compared to *T. diffusa* (C_3_) and *B. prostrata* (C_4_). This corroborates the expression patterns reported in the C_2_ species of *Flaveria* (Gowik et al., 2011; Mallmann et al., 2014) and *Salsoleae* (Lauterbach et al., 2017b), implying a successful integration of C_2_ photosynthesis in *Sed. sedoides* (C_2_). Our transcript data showed that GDC-P and a SHMT isoform (Bv6_143730_mggd.t1) were down-regulated in *Sed. sedoides* (C_2_) when compared to *T. diffusa* (C_3_) and *B. prostrata* (C_4_), respectively. Similar results were obtained in the C_2_ species of the genus *Flaveria* (Mallmann et al., 2014). Schulze et al. (2013) showed that down-regulation of GDC-P was closely linked to the establishment of the C_2_ pathway in *Flaveria*. Since GDC-P and a SHMT isoform are well known to be involved in glycine decarboxylation, their down-regulation in *Sed. sedoides* (C_2_) might have similar consequences. It is worth noticing that the number of significantly up-regulated photorespiratory genes in *Sed. sedoides* (C_2_) was equal to *T. diffusa* (C_3_) when compared to *B. prostrata* (C_4_).

A significant reduction of almost all photorespiratory genes was observable in *B. prostrata* (C_4_). All photorespiratory genes except GLYK were down-regulated in *B. prostrata* (C_4_) as compared to *T. diffusa* (C_3_). Mallmann et al. (2014) reported significant down regulation of all photorespiratory genes in C_4_ *Flaveria* except the transport proteins DIT1 and DIT2 and one isoform of GLDH. On the other hand, GLYK was found expressed in the M of C_4_ *Sorghum bicolor* (Döring *et al*., 2016). GLYK catalyses the regeneration of 3-phosphoglycerate (3-PG). The localisation of GLYK within the leaf cells of *B. prostrata* (C_4_) could clarify its high expression and role.

### Regulation of C_3_, C_2_ and C_4_ photosynthesis in Amaranthaceae/Chenopodiaceae

Transcription factors are proteins that bind to the DNA promoter or enhancer regions of specific genes and regulate their expression. They have a crucial role on plant growth, development and adaptation under various stress conditions, and therefore are excellent candidates for modifying complex traits in plants (Ambawat *et al*., 2013). C_3_, C_2_ and C_4_ species of *Salsoleae* and *Camphorosmeae* are widely spread in desert, semi-desert, saline, arid regions (Akhani *et al*., 2007; Kadereit et al., 2011). In former Chenopodiaceae, C_4_ photosynthesis is an adaptation to hot, dry or saline areas from C_3_ ancestor that was already preadapted to grow in these harsh environments (Kadereit *et al*., 2012). We focused on TFs that were differentially expressed between C_3_, C_2_, and C_4_ species/states irrespective of the lineage, to further reduce the amount of differentially expressed TFs to a small subset of actually C_4_, C_2_, C_3_-related changes. Indeed, a small number of TFs were found differentially expressed between C_3_, C_2_, and C_4_ species/states.

Cluster analysis showed that BBX15, SHR, SCZ and LBD41 were co-regulated and significantly higher abundant in all C_4_ species irrespective of the lineage when compared to C_3_ species. The families to which these TF families belong, play an important role in regulatory networks controlling plant growth and development, and plant adaptive responses to various environmental stress conditions (Kotak *et al*., 2004; Pernas *et al*., 2010; Schmidt *et al*., 2012; Gangappa and Botto, 2014; Grimplet *et al*., 2017). Except the LBD TF family, SHR, HSF, and CO-like have been shown to be involved in the development of C_4_ Kranz anatomy in *Zea mays* L. and potentially involved in the establishment of C_4_ M and Kranz cell identities (Slewinski *et al*., 2012; Slewinski, 2013; Wang *et al*., 2013; Fouracre *et al*., 2014; Lori Tausta *et al*., 2014). However, members of the LBD TF family are key regulators of plant organ development, leaf development, pollen development, plant regeneration, stress response, and anthocyanin and nitrogen metabolisms (Semiarti *et al*., 2001; Grimplet *et al*., 2017). Since mRNA of all the four TFs was highly abundant in C_4_ species and co-regulated, our data suggest a critical role of these TFs in the development of any C_4_ Kranz anatomy in the *Amaranthaceae*/*Chenopodiaceae* complex.

We found that C_3_ species enhanced different TFs compared to C_4_ species. Three TFs (ATHB13, HD-ZIP family), HSFA6B (HSF TF family), and NAC083 (NAC TF family), in which two TFs (HSF6B and NAC083) are co-regulated, were significantly higher in all C_3_ species when compared to C_4_ species. As C_4_ TFs, they are involved in plant growth, development, and stress tolerance. NAC TF family was shown to contribute to root and shoot apical meristems formation in *Arabidopsis* (Xie *et al*., 2000; Vroemen *et al*., 2003), organogenesis (Yamaguchi *et al*., 2010), salt and drought tolerance in *Arabidopsis* (Huang et al., 2015; Huang *et al*., 2015), leaf senescence in tobacco (Li *et al*., 2018) and secondary cell wall formation in cotton (Zhang et al., 2017; Li *et al*., 2018). HD-Zip TF family was reported to regulate plant growth adaptation to abiotic stress such as salt and drought in apple and *Arabidopsis* (Ebrahimian-Motlagh *et al*., 2017; Zhang *et al*., 2021). Interestingly, HD-ZIP, HSF and NAC TF families were suggested to control the C_4_ photosynthesis in maize and rice (Li *et al*., 2010; Lori Tausta *et al*., 2014). However, in these studies, these TF families were detected using development gradient transcriptome comparison only on C_4_ maize and rice plants. This may imply higher activity of these TF families in C_3_ species. Nevertheless, different transcripts of these TFs families were involved when compared to the present study. Thus, a significant expression of these TFs in C_3_ species could indicate a potential function of these TFs in the C_3_ pathway.

In C_2_ species, one transcript of the BASIC HELIX-LOOP-HELIX (bHLH106) protein from the bHLH TF family was found up-regulated as compared to C_4_ and C_3_ species. Two TFs of the bHLH TF family were shown to regulate a C_4_ photosynthesis gene in maize (Borba *et al*., 2018). This up-regulation of bHLH106 in all C_2_ species, may suggest its possible role in the development and establishment of the C_2_ photosynthesis specificities relative to other photosynthesis types. Interestingly, one C_4_ specific TF (BBX15, TF family CO-like) was significantly higher in C_2_ species when compared to C_3_ species. Thus, this TF could be responsible for similarities of C_4_ photosynthesis found in C_2_ species such as the Kranz-like anatomy. Surprisingly, no C_3_ specific TF was significantly expressed in C_2_ species when compared to C_4_ species. This indicates that C_2_ and C_4_ photosynthesis represent more derived types of photosynthesis compared to C_3_ photosynthesis. Nonetheless, this seems to be inconsistent with the current model of C_4_ evolution which relies heavily on the interpretation of the physiological intermediacy of C_2_ photosynthesis as an evolutionary stepping stone to C_4_ (Sage *et al*., 2018). One would expect if the C_2_ photosynthesis type represents an intermediate step along the C_3_-to-C_4_ photosynthesis that C_3_ specific TFs were somehow higher in C_2_ species when compared to C_4_ species as revealed by differential expression analysis of core photorespiratory genes in C_2_ and C_3_ species of *Camphorosmeae* (this study) and *Salsoleae* (Lauterbach et al., 2017b).

Taking the results of this study together, the unique derived TF profile of the C_2_ intermediate species suggest an evolutionary-stable state in its own right. Similarities with C_4_ relatives might result from a hybrid origin involving a C_3_ and a C_4_ parental lineage, parallel recruitment of a number of TFs in C_4_ and C_2_ lineages or a common ancestry and later divergent evolution. The position of *Sedobassia* as sister to *Bassia* (all C_4_) allows all of these three scenarios (Kadereit et al., 2014). For C_2_ species in *Salsoleae*, however, phylogenomic evidence points to a hybrid origin of the *Sal. divaricata* agg. (Tefarikis et al., submitted). If an early hybridization event of a C_4_ (or ancestral preadapted C_4_) lineage) and a C_3_ lineage led to the origin of the *Sedobassia* lineage which then evolved towards stable C_2_ photosynthesis awaits further phylogenomic analyses.

## Conclusion

Our transcriptome data of the *Chenopodiaceae* family provided new insight into C_4_ evolution. Proteins encoding for C_4_ transporters (DIT and TPT) were found significantly up- regulated in *Sed. sedoides* (C_2_) when compared to *B. prostrata* (C_4_). Up-regulation of those transporters reduces CO_2_ leakage from BSCs to MC, which could otherwise be detrimental to C_2_ photosynthesis due to the presence of RuBisCo in the MC. This suggests evolution of a stable C_2_ photosynthesis independent of C_4_ photosynthesis. Combined analysis of TFs of the sister lineages provided further support of this result. Indeed, while one C_4_ specific TF (BBX15) was significantly higher in C_2_ species when compared to C_3_ species, no C_3_ specific TFs were higher in C_2_ species as compared to C_4_ species. Finally, apart from well-known TFs involved in the development of C_4_ Kranz anatomy like SHR: BBX15, SCZ and LBD41 may as well associate with its development and physiology. Furthermore, bHLH106 could be related to specific C_2_ anatomy and BBX15 to a characteristic C_4_-like expression pattern found in species with C_2_ photosynthesis. This study sheds light on the differentiate regulation and evolution of transcription factors in C_2_ and C_4_ photosynthesis.

## Supplementary data

Table S1. Voucher information of species used in this study.

Dataset S1. Read Mapping statistics of *Camphorosmeae* species

Table S2. Quality assessment of transcriptome *de novo* assemblies using the Eudicotyledons odb10 dataset in BUSCO v.3.0. BUSCOs are categorized into complete, fragmented, and missing. Category ‘complete’ is subdivided into ‘complete and single-copy’ and ‘complete and duplicated’. The total number of BUSCOs in the Eudicotyledons odb10 dataset was 2121. B., *Bassia*; H., *Hammada*; Sal., *Salsola*; Sed., *Sedobassia*; T., *Threlkeldia*.

Dataset S2. Differential expression analysis of pairwise comparisons between *Camphorosmeae* species

Dataset S3. Transcriptional investment of *Camphorosmeae* species

Dataset S4. Annotation and normalized transcript count of C4-related and photorespiratory genes

Dataset S5. Clustered TFs

Fig. S1. All clusters of TFs per photosynthesis types

## Acknowledgements

We would like to thank Kumari Billakurthi and the late Udo Gowik for their useful advice and support during this study. We also thank the Millenium Seed Bank (MSB Kew), Jaime Gil González (Lanzarote), Magui Olangua Corral (Gran Canaria), Helmut Freitag (Kassel) and Elena Voznesenskaya (St. Petersburg) for the contribution of seed samples. We thank the “Genomics and Transcriptomics laboratory” of the “Biologisch-Medizinisches Forschungszentrum” (BMFZ) at the Heinrich-Heine-University Duesseldorf (Germany) for technical support and conducting the Illumina sequencing. Parts of this research were performed using the supercomputer Mogon and/or advisory services offered by Johannes Gutenberg University Mainz (hpc.uni-mainz.de), which is a member of the AHRP and the Gauss Alliance e.V. We thank C. Wild (Botanical Garden Univ. Mainz) for cultivating the plants.

## Author contributions

Conceptualization: GK; Data curation: ML, CS; Formal Analysis: ML, CS; Funding acquisition: GK; Investigation: ML; Methodology: GK; Project administration: GK; Resources: GK; Software: ML, CS; Supervision: GK; Validation: CS, ML; Visualization: CS, ML; Writing-original draft: CS, ML; Writing – review & editing: GK, CS.

## Funding

This work was funded by the Deutsche Forschungsgemeinschaft (DFG) with grants to GK (KA1816/7-3).

## Notes

### Competing Interest Statement

The authors have declared no competing interest.

## References

Abu-Jamous B, Kelly S. 2018. Clust: Automatic extraction of optimal co-expressed gene clusters from gene expression data. Clust: automatic extraction of optimal co-expressed gene clusters from gene expression data 19:172, 1–11.

Akhani H, Edwards G, Roalson EH. 2007. Diversification of the old world Salsoleae s.I. (Chenopodiaceae): Molecular phylogenetic analysis of nuclear and chloroplast data sets and a revised classification. International Journal of Plant Sciences 168, 931–956.

Ambawat S, Sharma P, Yadav NR, Yadav RC. 2013. MYB transcription factor genes as regulators for plant responses: An overview. Physiology and Molecular Biology of Plants 19, 307–321.

Aubry S, Kelly S, Kümpers BMC, Smith-Unna RD, Hibberd JM. 2014. Deep Evolutionary Comparison of Gene Expression Identifies Parallel Recruitment of Trans-Factors in Two Independent Origins of C4 Photosynthesis. PLoS Genetics 10.

Bauwe H, Hagemann M, Fernie AR. 2010. Photorespiration: players, partners and origin. Trends in Plant Science 15, 330–336.

Bolger AM, Lohse M, Usadel B. 2014. Trimmomatic: A flexible trimmer for Illumina sequence data. Bioinformatics 30, 2114–2120.

Borba AR, Serra TS, Górska A, et al. 2018. Synergistic binding of bHLH transcription factors to the promoter of the maize NADP-ME gene used in C4 photosynthesis is based on an ancient code found in the ancestral C3 state. Molecular Biology and Evolution 35, 1690–1705.

Bräutigam A, Gowik U. 2016. Photorespiration connects C3 and C4 photosynthesis. Journal of Experimental Botany 67, 2953–2962.

Bräutigam A, Kajala K, Wullenweber J, et al. 2011. An mRNA blueprint for C4 photosynthesis derived from comparative transcriptomics of closely related C3 and C4 species. Plant Physiology 155, 142–156.

Christin PA, Osborne CP, Chatelet DS, Columbus JT, Besnard G, Hodkinson TR, Garrison LM, Vorontsova MS, Edwards EJ. 2013. Anatomical enablers and the evolution of C4 photosynthesis in grasses. Proceedings of the National Academy of Sciences of the United States of America 110, 1381–1386.

Dohm JC, Minoche AE, Holtgräwe D, et al. 2014. The genome of the recently domesticated crop plant sugar beet (Beta vulgaris). Nature 505, 546–549.

Döring F, Streubel M, Bräutigam A, Gowik U. 2016. Most photorespiratory genes are preferentially expressed in the bundle sheath cells of the C4 grass Sorghum bicolor. Journal of Experimental Botany 67, 3053–3064.

Ebrahimian-Motlagh S, Ribone PA, Thirumalaikumar VP, Allu AD, Chan RL, Mueller-Roeber B, Balazadeh S. 2017. JUNGBRUNNEN1 confers drought tolerance downstream of the HD-Zip I Transcription factor AtHB13. Frontiers in Plant Science 8, 1–12.

Edwards GE, Ku MSB. 1987. Biochemistry of C3–C4 Intermediates. ACADEMIC PRESS, INC.

Fouracre JP, Ando S, Langdale JA. 2014. Cracking the Kranz enigma with systems biology. Journal of Experimental Botany 65, 3327–3339.

Freitag H, Kadereit G. 2014. C3 and C4 leaf anatomy types in Camphorosmeae (Camphorosmoideae, Chenopodiaceae). Plant Systematics and Evolution 300, 665–687.

Fu L, Niu B, Zhu Z, Wu S, Li W. 2012. CD-HIT: Accelerated for clustering the next-generation sequencing data. Bioinformatics 28, 3150–3152.

Furumoto T, Yamaguchi T, Ohshima-Ichie Y, et al. 2011. A plastidial sodium-dependent pyruvate transporter. Nature 476, 472–476.

Kadereit G, Borsch T, Weising K and Freitag H. 2003. Phylogeny of Amaranthaceae and Chenopodiaceae and the evolution of C4 photosynthesis. Int. J. Plant Sci. 164, 959–986.

Gangappa SN, Botto JF. 2014. The BBX family of plant transcription factors. Trends in Plant Science 19, 460–470.

Gowik U, Bräutigam A, Weber KL, Weber APM, Westhoff P. 2011. Evolution of C 4 photosynthesis in the genus flaveria: How many and which genes does it take to make C 4? Plant Cell 23, 2087–2105.

Gowik U, Westhoff P. 2011. The Path from C3 to C4 photosynthesis. Plant Physiology 155, 56– 63.

Grabherr MG., Brian J. Haas, Yassour M et al. 2013. Trinity: reconstructing a full-length transcriptome without a genome from RNA-Seq data. Nature Biotechnology 29, 644–652.

Grimplet J, Pimentel D, Agudelo-Romero P, Martinez-Zapater JM, Fortes AM. 2017. The LATERAL ORGAN BOUNDARIES Domain gene family in grapevine: Genome-wide characterization and expression analyses during developmental processes and stress responses. Scientific Reports 7, 1–18.

Huang P, Brutnell TP. 2016. A synthesis of transcriptomic surveys to dissect the genetic basis of C4 photosynthesis. Current Opinion in Plant Biology 31, 91–99.

Huang Q, Wang Y, Li B, Chang J, Chen M, Li K, Yang G, He G. 2015. TaNAC29, a NAC transcription factor from wheat, enhances salt and drought tolerance in transgenic Arabidopsis. BMC Plant Biology 15, 1–15.

Jin J, Tian F, Yang DC, Meng YQ, Kong L, Luo J, Gao G. 2017. PlantTFDB 4.0: Toward a central hub for transcription factors and regulatory interactions in plants. Nucleic Acids Research 45, D1040–D1045.

Jin J, Zhang H, Kong L, Gao G, Luo J. 2014. PlantTFDB 3.0: A portal for the functional and evolutionary study of plant transcription factors. Nucleic Acids Research 42, 1182–1187.

Kadereit G, Ackerly D, Pirie MD. 2012. A broader model for C4 photosynthesis evolution in plants inferred from the goosefoot family (chenopodiaceae s.s.). Proceedings of the Royal Society B: Biological Sciences 279, 3304–3311.

Kadereit G, Bohley K, Lauterbach M, Tefarikis DT, Kadereit JW. 2017. C3–C4 intermediates may be of hybrid origin – a reminder. New Phytologist 215, 70–76.

Kadereit G, Freitag H. 2011. Molecular phylogeny of Camphorosmeae (Camphorosmoideae, Chenopodiaceae): Implications for biogeography, evolution of C4-photosynthesis and taxonomy. Taxon 60, 51–78.

Kadereit G, Lauterbach M, Pirie MD, Arafeh R, Freitag H. 2014. When do different C4 leaf anatomies indicate independent C 4 origins? Parallel evolution of C4 leaf types in Camphorosmeae (Chenopodiaceae). Journal of Experimental Botany 65, 3499–3511.

Kotak S, Port M, Ganguli A, Bicker F, Von Koskull-Döring P. 2004. Characterization of C-terminal domains of Arabidopsis heat stress transcription factors (Hsfs) and identification of a new signature combination of plant class a Hsfs with AHA and NES motifs essential for activator function and intracellular localization. Plant Journal 39, 98–112.

Kriventseva E V., Kuznetsov D, Tegenfeldt F, Manni M, Dias R, Simão FA, Zdobnov EM. 2019. OrthoDB v10: Sampling the diversity of animal, plant, fungal, protist, bacterial and viral genomes for evolutionary and functional annotations of orthologs. Nucleic Acids Research 47, D807–D811.

Langmead B, Salzberg SL. 2012. Fast gapped-read alignment with Bowtie 2. Nature Methods 9, 357–359.

Lauterbach M, Billakurthi K, Kadereit G, Ludwig M, Westhoff P, Gowik U. 2017a. C3 cotyledons are followed by C4 leaves: Intra-individual transcriptome analysis of Salsola soda (Chenopodiaceae). Journal of Experimental Botany 68, 161–176.

Lauterbach M, Schmidt H, Billakurthi K, Hankeln T, Westhoff P, Gowik U, Kadereit G. 2017b. De novo transcriptome assembly and comparison of C3, C3-C4, and C4 species of tribe salsoleae (Chenopodiaceae). Frontiers in Plant Science 8.

Li W, Godzik A. 2006. Cd-hit: A fast program for clustering and comparing large sets of protein or nucleotide sequences. Bioinformatics 22, 1658–1659.

Li H, Handsaker B, Wysoker A, Fennell T, Ruan J, Homer N, Marth G, Abecasis G, Durbin R. 2009. The Sequence Alignment/Map format and SAMtools. Bioinformatics 25, 2078–2079.

Li W, Li X, Chao J, Zhang Z, Wang W, Guo Y. 2018. NAC family transcription factors in tobacco and their potential role in regulating leaf senescence. Frontiers in Plant Science 871, 1– 15.

Li P, Ponnala L, Gandotra N, et al. 2010. The developmental dynamics of the maize leaf transcriptome. Nature Genetics 42, 1060–1067.

Lori Tausta S, Li P, Si Y, Gandotra N, Liu P, Sun Q, Brutnell TP, Nelson T. 2014. Developmental dynamics of Kranz cell transcriptional specificity in maize leaf reveals early onset of C4-related processes. Journal of Experimental Botany 65, 3543–3555.

Lundgren MR. 2020. C2 photosynthesis: a promising route towards crop improvement? New Phytologist 228, 1734–1740.

Mallmann J, Heckmann D, Bräutigam A, Lercher MJ, Weber APM, Westhoff P, Gowik U. 2014. The role of photorespiration during the evolution of C4 photosynthesis in the genus Flaveria. eLife 2014.

Monson RK, Edwards GE, Ku MSB. 1984. C3-C4 Intermediate Photosynthesi Plants. BioScience 34, 563–566.

Oakley JC, Sultmanis S, Stinson CR, Sage TL, Sage RF. 2014. Comparative studies of C3 and C4 Atriplex hybrids in the genomics era: Physiological assessments. Journal of Experimental Botany 65, 3637–3647.

Pernas M, Ryan E, Dolan L. 2010. SCHIZORIZA Controls Tissue System Complexity in Plants. Current Biology 20, 818–823.

Pyankov VI, Artyusheva EG, Edwards GE, Black CC, Soltis PS. 2001. Phylogenetic analysis of tribe salsoleae (Chenopodiaceae) based on ribosomal its sequences: Implications for the evolutIon of photosynthesis types. American Journal of Botany 88, 1189–1198.

Robinson MD, McCarthy DJ, Smyth GK. 2010. edgeR: A Bioconductor package for differential expression analysis of digital gene expression data. Bioinformatics 26, 139–140.

Sage RF. 2016. A portrait of the C4 photosynthetic family on the 50th anniversary of its discovery: Species number, evolutionary lineages, and Hall of Fame. Journal of Experimental Botany 67, 4039–4056.

Sage RF. 2021. Russ Monson and the evolution of C4 photosynthesis. Oecologia.

Sage TL, Busch FA, Johnson DC, Friesen PC, Stinson CR, Stata M, Sultmanis S, Rahman BA, Rawsthorne S, Sage RF. 2013. Initial events during the evolution of C4 photosynthesis in C3 species of Flaveria. Plant Physiology 163, 1266–1276.

Sage RF, Khoshravesh R, Sage TL. 2014. From proto-Kranz to C4 Kranz: Building the bridge to C 4 photosynthesis. Journal of Experimental Botany 65, 3341–3356.

Sage RF, Monson RK, Ehleringer JR, Adachi S, Pearcy RW. 2018. Some like it hot: the physiological ecology of C4 plant evolution. Oecologia 187, 941–966.

Sage RF, Sage TL, Pearcy RW, Borsch T. 2007. The taxonomic distribution of C4 photosynthesis in Amaranthaceae sensu stricto. American Journal of Botany 94, 1992–2003.

Schlüter U, Weber APM. 2020. Regulation and Evolution of C 4 Photosynthesis., 183–215.

Schmidt R, Schippers JHM, Welker A, Mieulet D, Guiderdoni E, Mueller-Roeber B. 2012. Transcription factor oshsfc1b regulates salt tolerance and development in oryza sativa ssp. japonica. AoB PLANTS 12.

Schulze S, Mallmann J, Burscheidt J, Koczor M, Streubel M, Bauwe H, Gowik U, Westhoff P. 2013. Evolution of C4 photosynthesis in the genus flaveria: Establishment of a photorespiratory CO2 pump. Plant Cell 25, 2522–2535.

Schüssler C, Freitag H, Koteyeva N, Schmidt D, Edwards G, Voznesenskaya E, Kadereit G. 2017. Molecular phylogeny and forms of photosynthesis in tribe Salsoleae (Chenopodiaceae). Journal of Experimental Botany 68, 207–223.

Schütze P, Freitag H, Weising K. 2003. An integrated molecular and morphological study of the subfamily Suaedoideae Ulbr. (Chenopodiaceae). Plant Systematics and Evolution 239, 257– 286.

Schwacke R, Ponce-Soto GY, Krause K, Bolger AM, Arsova B, Hallab A, Gruden K, Stitt M, Bolger ME, Usadel B. 2019. MapMan4: A Refined Protein Classification and Annotation Framework Applicable to Multi-Omics Data Analysis. Molecular Plant 12, 879–892.

Semiarti E, Ueno Y, Tsukaya H, Iwakawa H, Machida C. 2001. The ASYMMETRIC LEAVES2 gene of Arabidopsis thaliana regulates formation of a symmetric lamina, establishment of venation and repression of meristem-related homeobox genes in leaves. 1783, 1771–1783.

Simão FA, Waterhouse RM, Ioannidis P, Kriventseva E V., Zdobnov EM. 2015. BUSCO: Assessing genome assembly and annotation completeness with single-copy orthologs. Bioinformatics 31, 3210–3212.

Slewinski TL. 2013. Using evolution as a guide to engineer Kranz-type C4 photosynthesis. Frontiers in Plant Science 4, 1–13.

Slewinski TL, Anderson AA, Zhang C, Turgeon R. 2012. Scarecrow plays a role in establishing Kranz anatomy in maize leaves. Plant and Cell Physiology 53, 2030–2037.

Voznesenskaya E V., Koteyeva NK, Akhani H, Roalson EH, Edwards GE. 2013. Structural and physiological analyses in Salsoleae (Chenopodiaceae) indicate multiple transitions among C3, intermediate, and C4 photosynthesis. Journal of Experimental Botany 64, 3583–3604.

Vroemen CW, Mordhorst AP, Albrecht C, Kwaaitaal MACJ, De Vries SC. 2003. The CUP-SHAPED COTYLEDON3 gene is required for boundary and shoot meristem formation in Arabidopsis. Plant Cell 15, 1563–1577.

Walters RG, Ibrahim DG, Horton P, Kruger NJ. 2004. A mutant of arabidopsis lacking the triose-phosphate/phosphate translocator reveals metabolic regulation of starch breakdown in the light. Plant Physiology 135, 891–906.

Wang Y, Bräutigam A, Weber APM, Zhu XG. 2014. Three distinct biochemical subtypes of C4 photosynthesis? A modelling analysis. Journal of Experimental Botany 65, 3567–3578.

Wang P, Kelly S, Fouracre JP, Langdale JA. 2013. Genome-wide transcript analysis of early maize leaf development reveals gene cohorts associated with the differentiation of C4 Kranz anatomy. Plant Journal 75, 656–670.

Windhövel A, Hein I, Dabrowa R, Stockhaus J. 2001. Characterization of a novel class of plant homeodomain proteins that bind to the C4 phosphoenolpyruvate carboxylase gene of Flaveria trinervia. Plant Molecular Biology 45, 201–214.

Xie Q, Frugis G, Colgan D, Chua NH. 2000. Arabidopsis NAC1 transduces auxin signal downstream of TIR1 to promote lateral root development. Genes and Development 14, 3024– 3036.

Yamaguchi M, Ohtani M, Mitsuda N, Kubo M, Ohme-Takagi M, Fukuda H, Demura T. 2010. VND-INTERACTING2, a NAC domain transcription factor, negatively regulates xylem vessel formation in Arabidopsis. Plant Cell 22, 1249–1263.

Zhang Q, Chen T, Wang X, Wang J, Gu K, Yu J, Hu D, Hao Y. 2021. Genome-wide Identification and Expression Analyses of Homeodomain-leucine Zipper Family Genes Reveal Their Involvement in Stress Response in Apple (Malus × domestica). Horticultural Plant Journal.

