## Supplementary figures and images for "Insights into regulation of C_2_ and C_4_ photosynthesis in *Amaranthaceae/Chenopodiaceae* using RNA-Seq"

### Figure S1

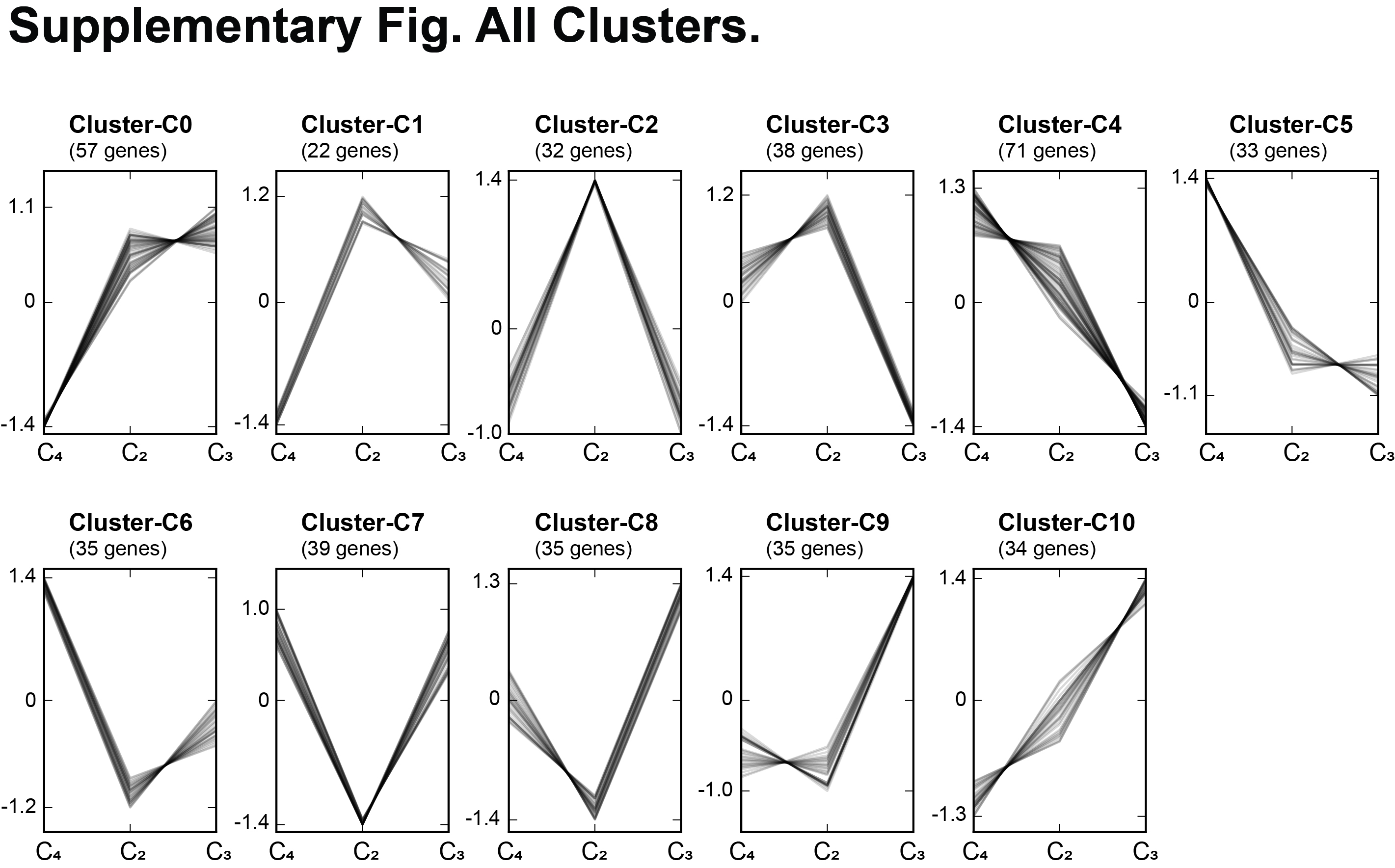
