## Supplementary material for "Insights into regulation of C_2_ and C_4_ photosynthesis in *Amaranthaceae/Chenopodiaceae* using RNA-Seq": Table S1

**Table Voucher**.

| Lineage | Species | PS type | Herbarium/voucher ID |
| --- | --- | --- | --- |
| Camphorosmeae | ***Threlkeldia diffusa*** R. Br. | C_3_ | Millenium Seed Bank serial number: 0381990; cultivated at Botanical Garden Mainz, living coll. no 076 (voucher MJG 027679) |
|  | ***Sedobassia sedoides*** (Pall.) Freitag & G.Kadereit | C_2_ | G. Somogyi s.n., Hungary, Besenyötelek, Csuportos-legelö, N 47,688119°, E 20,461372°, 2012 (cultivated at Botanical Garden Mainz, living coll. no 002; voucher at MJG) |
|  | ***Bassia prostrata*** (L.) Beck | C_4_ | G. Somogyi s.n., Hungary, Besenyötelek, Csuportos-legelö, N 47,685817°, E20,465919°, 2012 (cultivated at Botanical Garden Mainz, living coll. no 001; voucher at MJG) |
| Salsoleae | ***Salsola webbii*** Moq. | C_3_ | Elena Voznesenskaya s.n., Southern Spain, collected in 2011; (cultivated at Pullman Univ., Washington and Botanical Garden Mainz; living coll. number 67; voucher at MJG) |
|  | ***Salsola divaricata*** Masson ex Link  Pop-184 | C_2_ | Gil González, J. s.n., 2013-10-21, Spain, Canary Islands, Lanzarote, Haria, Punta Mujeres (MJG 014225) |
|  | ***Salsola divaricata*** Masson ex Link  Pop-198 | C_2_ | Anonymous collector, Spain, Gran Canaria, Cuesta Ramón, Jinamar, Zona 28R, UTM: E 458720,932 N 3102402,105 (cultivated at Botanical Garden Mainz, living collection no 198II; MJG 013542) |
|  | ***Salsola soda*** L. | C_4_* | Kaligarič, M. s.n. 2009-09-14, Slovenia, Sv. Katarina (Ankaran) near Koper (cultivated at Botanical Garden Mainz living collection no. 125; MJG 013542) |
|  | *Hammada scoparia (Pomel) Iljin synonym of*  ***Haloxylon scoparium*** *Pomel* | C_4_ | Millenium Seed Bank serial number 0089920, cultivated at Botanical Garden Mainz, living coll. no 087 and 089. |
|  | ***Salsola oppositifolia*** Desf. | C_4_ | H. Freitag 40002; SW Morocco 18 km N Agadir, near Tamrhakt, behind coastal cliff, most common along roadside (cultivated at Botanical Garden Mainz, living coll. number 173; MJG 013564) |
