## Supplementary material for "Insights into regulation of C_2_ and C_4_ photosynthesis in *Amaranthaceae/Chenopodiaceae* using RNA-Seq": Table S2

**Supplementary Table BUSCO.** Quality assessment of transcriptome *de novo* assemblies using the Eudicotyledons odb10 dataset in BUSCO v.3.0. BUSCOs are categorized into complete, fragmented, and missing. Category ‘complete’ is subdivided into ‘complete and single-copy’ and ‘complete and duplicated’. The total number of BUSCOs in the Eudicotyledons odb10 dataset was 2121. B., *Bassia*; H., *Hammada*; Sal., *Salsola*; Sed., *Sedobassia*; T., *Threlkeldia*.

| **Lineage** | **Assembly** | **Complete** | | **Complete, single** | | **Complete, duplicated** | | **Fragmented** | | **Missing** | |
| --- | --- | --- | --- | --- | --- | --- | --- | --- | --- | --- | --- |
|  |  | **Total** | **%** | **Total** | **%** | **Total** | **%** | **Total** | **%** | **Total** | **%** |
| Camphorosmeae | *B. prostrata* | 1879 | 88.6 | 864 | 40.7 | 1015 | 47.9 | 121 | 5.7 | 121 | 5.7 |
|  | *Sed. sedoides* | 1929 | 90.9 | 1148 | 54.1 | 781 | 36.8 | 97 | 4.6 | 95 | 4.5 |
|  | *T. diffusa* | 1868 | 88.1 | 965 | 45.5 | 903 | 42.6 | 142 | 6.7 | 111 | 5.2 |
| Salsoleae | *H. scoparia* | 1818 | 85.7 | 1089 | 51.3 | 729 | 34.4 | 183 | 8.6 | 120 | 5.7 |
|  | *Sal. divaricata 184* | 1846 | 87.0 | 599 | 28.2 | 1247 | 58.8 | 164 | 7.7 | 111 | 5.3 |
|  | *Sal. divaricata 198* | 1832 | 86.4 | 606 | 28.6 | 1226 | 57.8 | 190 | 9.0 | 99 | 4.6 |
|  | *Sal. oppositifolia* | 1733 | 81.7 | 480 | 22.6 | 1253 | 59.1 | 255 | 12.0 | 133 | 6.3 |
|  | *Sal. soda* | 1914 | 90.2 | 1273 | 60.0 | 641 | 30.2 | 126 | 5.9 | 81 | 3.9 |
|  | *Sal. webbii* | 1814 | 85.5 | 585 | 27.6 | 1229 | 57.9 | 149 | 7.0 | 158 | 7.5 |
